# Metastatic function of METTL18 in breast cancer via actin methylation and Src

**DOI:** 10.1101/831701

**Authors:** Han Gyung Kim, Ji Hye Kim, Woo Seok Yang, Jae Gwang Park, Yong Gyu Lee, Yo Han Hong, Eunji Kim, Minkyeong Jo, Chae Young Lee, Shi Hyung Kim, Nak Yoon Sung, Young-Su Yi, Zubair Ahmed Ratan, Sunggyu Kim, Byong Chul Yoo, Sung-Ung Kang, Young Bong Kim, Sangmin Kim, Hyun-June Paik, Jeong Eon Lee, Seok Jin Nam, Narayanan Parameswaran, Jeung-Whan Han, Jae Youl Cho

## Abstract

Recently, a SET domain containing 3 (SETD3) was identified as an actin histidine methyltransferase, functioning to control replication and pathogenesis in multiple mouse models for enterovirus infection as well as the regulation of smooth muscle contractility linked to primary dystocia. Here, in this study, we report another type of actin histidine methyltransferase, METTL18, that regulates the metastatic potential of breast cancer in human. Among methyltransferases, METTL18 was highly amplified in human breast cancer. In particular, poor prognosis was associated with high expression of METTL18 in HER2-negative breast cancer patients. This gene product was also found to be a critical component of metastatic responses. Loss of METTL18 expression significantly reduced metastatic responses of breast tumor cells both *in vitro* and *in vivo*. Mechanistically, it was observed that METTL18 increased actin polymerization, upregulated complex formation with HSP90AA1 and Src, enhanced the activity of an intermediate form of Src with tyrosine phosphorylation at both Y416 and Y527, and induced cellular metastatic responses, including morphological change, migration, and invasion of MDA-MB-231 cells *in vitro* and in mice. Methylated actin at His73 served as a critical site for interaction with HSP90AA1 and Src to activate p85/PI3K and STAT3. Our findings suggest that METTL18 plays critical roles in metastatic responses of HER2-negative breast cancer cells via actin polymerization and the generation of an intermediate form of Src.

## Introduction

Breast cancer is the most common form of malignant tumor and the second leading cause of cancer-related deaths in women (Siegel et al, 2018). A heterogeneous disease, breast cancer is classified into four intrinsic subtypes depending on the presence of estrogen receptor (ER), progesterone receptor (PR), and human epidermal receptor 2 (HER2) biomarkers: (1) luminal A: HER2-negative; (2) luminal B: ER-positive; (3) HER2 overexpression: HER2-positive; and 4) triple negative breast cancer (TNBC): HER2, ER, and PR negative (Aysola et al, 2013; Gao & Swain, 2018; Vallejos et al, 2010). Of these, luminal A with the HER2-negative phenotype is the most common subtype, accounting for 54.3% of all breast cancer patients (O’Brien et al, 2010) and known to have higher metastatic potential to bone (Kennecke et al, 2010). TNBC, another subtype without HER2, accounts for 10–15% of breast cancers and is more difficult to diagnose, more aggressive, and has a higher recurrence rate than other hormone receptor-positive breast cancers (Lebert et al, 2018; Waks & Winer, 2019). In addition, there are no approved drugs targeting TNBC at present and optimal chemotherapy has not yet been established (Lebert et al, 2018). Nonetheless, the motility of breast cancer is critically decided by the metastatic potential of these cells, with metastatic breast cancer accounting for up to 80% of breast cancer deaths compared to 10% for localized carcinoma (Mukherjee & Zhao, 2013). Therefore, it is necessary to explore molecules that can be potential therapeutic targets based on the present knowledge of tumorigenesis in TNBC as well as metastatic events of breast cancer cells. In addition, diverse types of breast cancer proceed through different mechanisms, and these differences need to be understood for accurate diagnosis and treatment.

METTL18 (a histidine methyltransferase 1 homolog, also known as C1orf156 or arsenic-transactivated protein 2, AsTPs) is a protein belonging to the methyltransferase-like protein (METTL) family (http://www.genenames.org/cgi-bin/genefamilies/set/963) and is considered a putative human homolog of YY110W, yeast histidine methyltransferase (Webb et al, 2010). In nature, only a few proteins, including β-actin (Johnson et al, 1967), S100A9 (Raftery et al, 1996), myosin (Elzinga & Collins, 1977), myosin kinase (Meyer & Mayr, 1987), and ribosomal protein RPL3 (Webb et al, 2010), are known to be methylated at histidine residues. Interestingly, a recent study revealed that protein methylation on histidine residues may be quite common in intracellular proteins in mammalian cells (Lappalainen, 2019). However, research in this area is difficult due to the lack of effective reagents for probing protein-histidine methylation (Ning et al, 2016). Recently, SET domain containing 3 (SETD3), which was previously shown to act as a histone lysine methyltransferase, was identified as an actin histidine methyltransferase, functioning to control replication and pathogenesis in multiple mouse models for enterovirus infection as well as the regulation of smooth muscle contractility linked to primary dystocia (Diep et al, 2019; Kwiatkowski et al, 2018; Wilkinson et al, 2019). In contrast, however, the biological function and molecular characterization of METTL18 have yet to be elucidated.

Tyrosine kinases, highly expressed in numerous types of tumor cells, are important molecules that manage the signal transduction process, morphological change, migration, metastasis, proliferation, metabolism, and programmed cell death (Paul & Mukhopadhyay, 2004). Src is a tyrosine kinase that has been recognized as a multiple player in tumorigenic responses and has been observed in about half the tumors from lung, brain, stomach, liver, colon, and breast cancers (Dehm & Bonham, 2004; Skorski, 2002). Interestingly, adenosine dialdehyde (AdOx), a methylation inhibitor (Bartel & Borchardt, 1984; Hong et al, 2008), can control metastatic responses of breast tumor cells such as morphological changes, migration, and invasion, and actin cytoskeleton disruption and the blockade of Src kinase activity are exhibited in AdOx treatment (Kim et al, 2013). Because AdOx is a transmethylation inhibitor, a methyltransferase that could methylate actin was predicted as a target enzyme in the AdOx-induced suppression of Src activity and metastatic potential (Kim et al, 2013). Based on this background, we examined methylation of the histidine (His)-73 residue of actin by METTL18 as a crucial component of metastatic responses of breast tumor cells through modulation of the actin cytoskeleton and Src phosphorylation. Since SETD3 is the first known actin histidine methyltransferase, we also compared molecular and cellular features between SETD3 and METTL18 in breast cancer cells.

## Results

### Clinical relevance of METTL18 in human breast cancer

To understand the biological role of METTL18 in metastatic responses, we studied its function in the context of breast tumors. To examine gene expression in breast cancer patients, we analyzed the mRNA expression level of METTL18 with other 25 methyltransferases in 442 patients from the TCGA data set that was classified according to PAM50 subtype of breast cancer (Fig. 1a). The expression level of METTL18 mRNA was highly amplified in breast tumors and especially highly expressed in HER2-negative patients, although the top gene was identified as SETDB1 (Fig. 1a). To determine whether similar mRNA expression was also present in Korean breast cancer patients, mRNA and protein expression levels of METTL18 in tissue samples from 150 Korean breast cancer patients were evaluated. The expression level of METTL18 was higher in two HER2-negative groups (Luminal A and TNBC stages) but not in two HER2-positive groups (Luminal B and HER2-postive stages) (Fig. 1b and Supplementary Fig. S1). Of these patients, those with higher METTL18 levels experienced significantly lower overall survival (p=0.037) (Fig. 1c). Similarly, Kaplan-Meier plots indicated that HER2-negative patients with higher METTL18 levels had a lower probability of survival (Fig. 1d), suggesting that METTL18 is positively involved in cancerous responses.

**Fig. 1.**
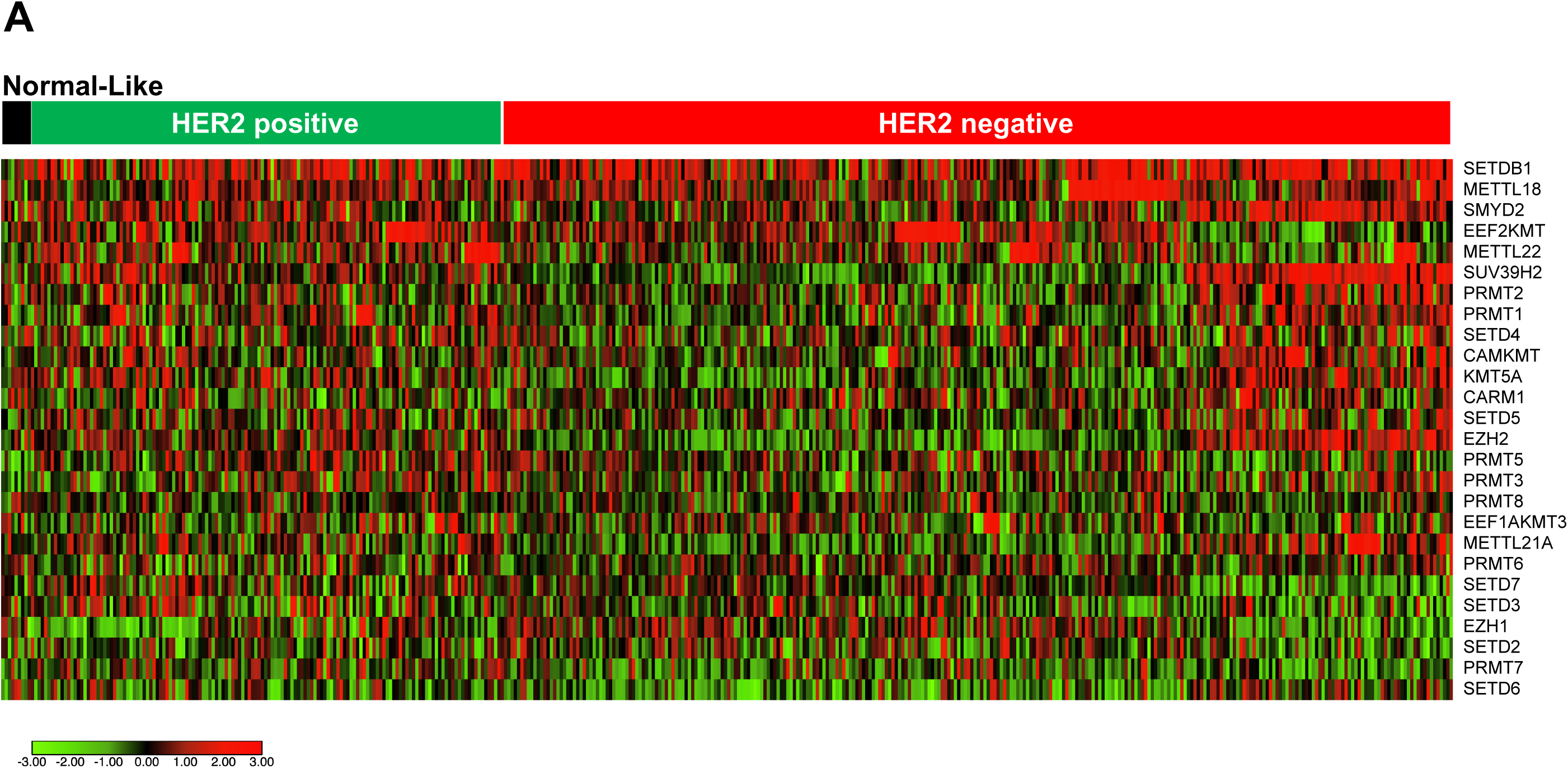
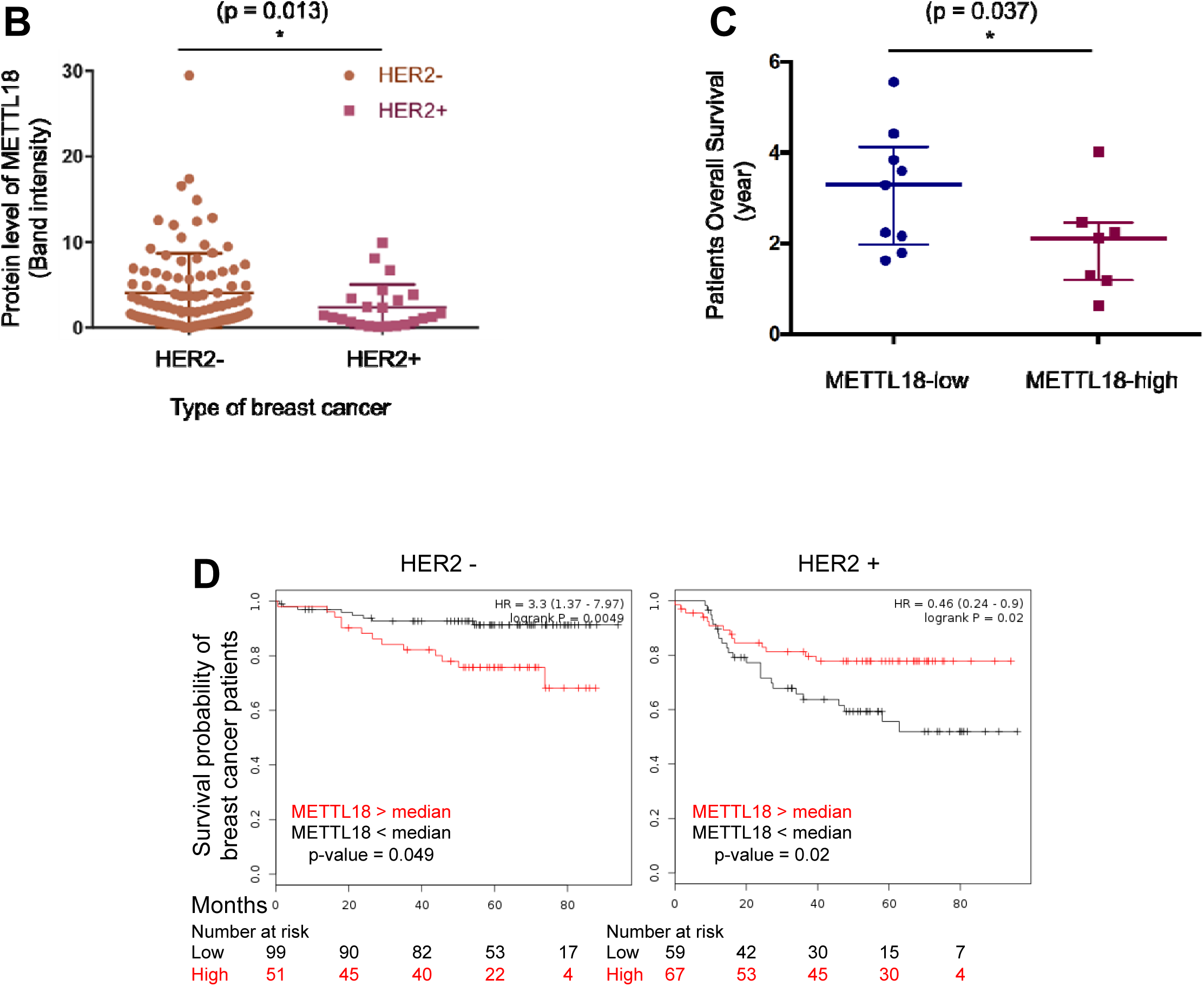
Clinical relevance of METTL18 in human breast cancer. (**A**) Heatmap of METTL18 and 25 methyltransferases expression profiles used with the TCGA breast cancer dataset. Heatmap was analyzed using www.software.broadinginstitute.org. (**B**) Protein levels of METTL18 from HER2 negative and positive groups were examined by Western blotting and semi-quantitative polymerase chain reaction (PCR). (**C**) Overall survival year of breast cancer patients with low or high levels of METTL18 was determined by survival time of the patient. Levels of METTL18 were measured by Western blotting analysis. (**D**) Kaplan-Meier curve of the relationship between survival probability and time (Month) was based on METTL18 expression levels from breast cancer patients with HER2-negative and - positive stages (*: p<0.05).

### Effect of METTL18 on metastatic capacity of breast cancer cells *in vitro* and *in vivo*

To examine whether METTL18 plays a critical role in metastatic responses *in vitro* and *in vivo*, we first tested the expression level of METTL18 in breast cancer cell lines. Protein levels of METTL18 were higher in HER2-negative cell lines such as MDA-MB-231 (TNBC cell line) and MCF-7 (Luminal A type breast cancer cell line), while HER2-enriched cell lines such as MDA-MB-453 and SK-BR3 showed low METTL18 expression (Fig. 2a). We next further established a knockdown cell line using MDA-MB-231 cells with TNBC properties and without HER2 expression (Chavez et al, 2010). Two different METTL18 knockdown cell lines (#2975 and #2974) were established (using two different shRNAs against METTL18) (Supplementary Fig. S2a). The migration and invasion of MDA-MB-231 cells were severely affected by both knockdown cell lines with shRNAs (#2975 and #2974) and overexpression of METTL18 (Fig. 2b-c and Supplementary Fig. S2b-c). The activity of matrix degradable enzymes (MMP-9 and MMP-2) was also suppressed through knockdown of METTL18 (Fig. 2d and Supplementary Fig. S2d and S2e) (van Kempen & Coussens, 2002). Together, these *in vitro* experiments demonstrated a critical role for METTL18 in metastasis of breast cancer cells. We next assessed the role of METTL18 in metastasis *in vivo*. To achieve this, MDA-MB-231 cells expressing shMETTL18 (#2975) were intravenously injected into nude mice to construct xenograft and metastatic models. These mice were then followed for four weeks using ^18^F-FDG-PET/CT scans. Consistent with our *in vitro* findings, metastatic dispersion of MDA-MB-213 cells in the nude mice was significantly reduced (p<0.005) with METTL18 knockdown cells compared to control cells (Fig. 2e). Body weight, however, did not differ between the groups (Supplementary Fig. S3).

**Fig. 2.**
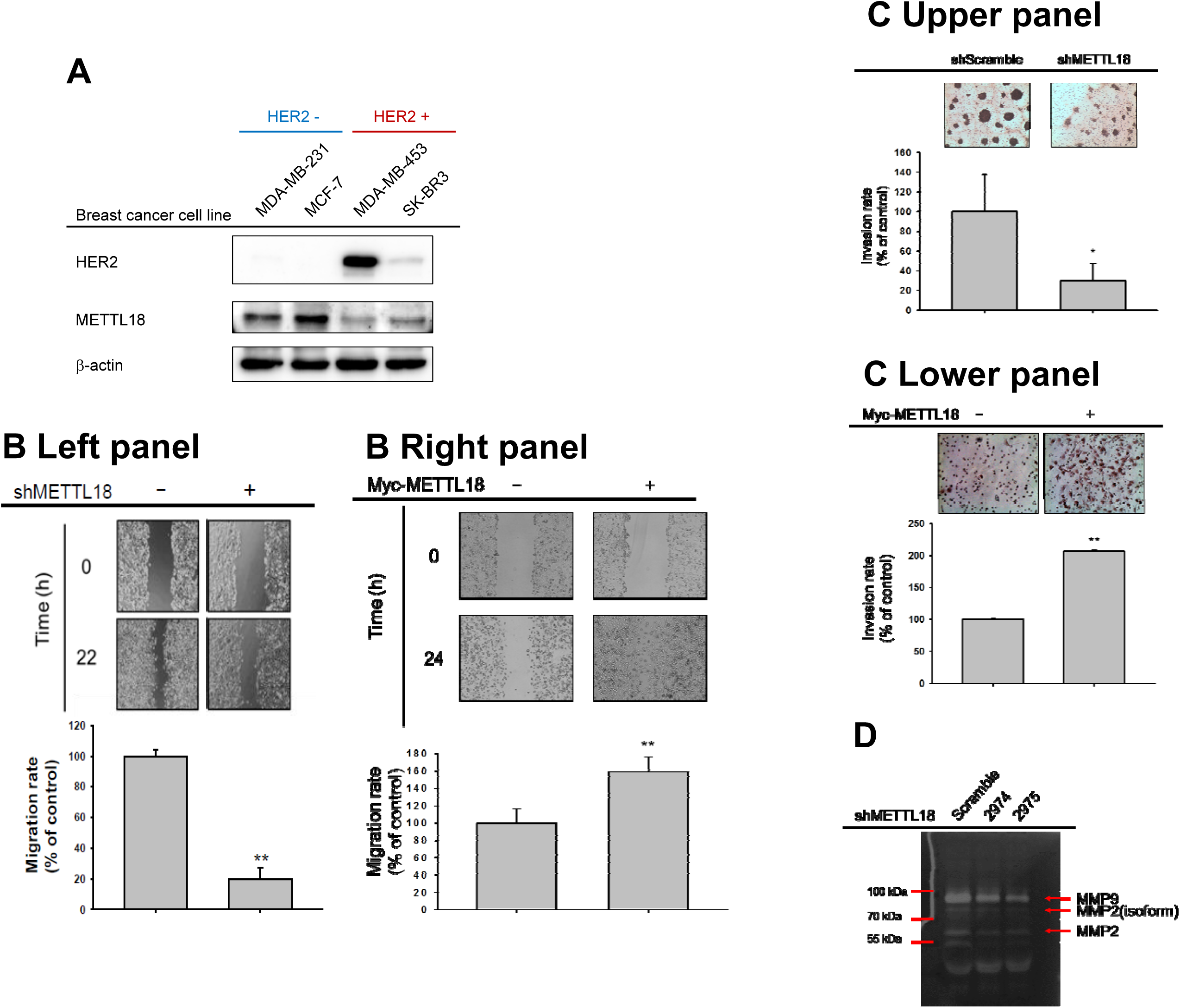
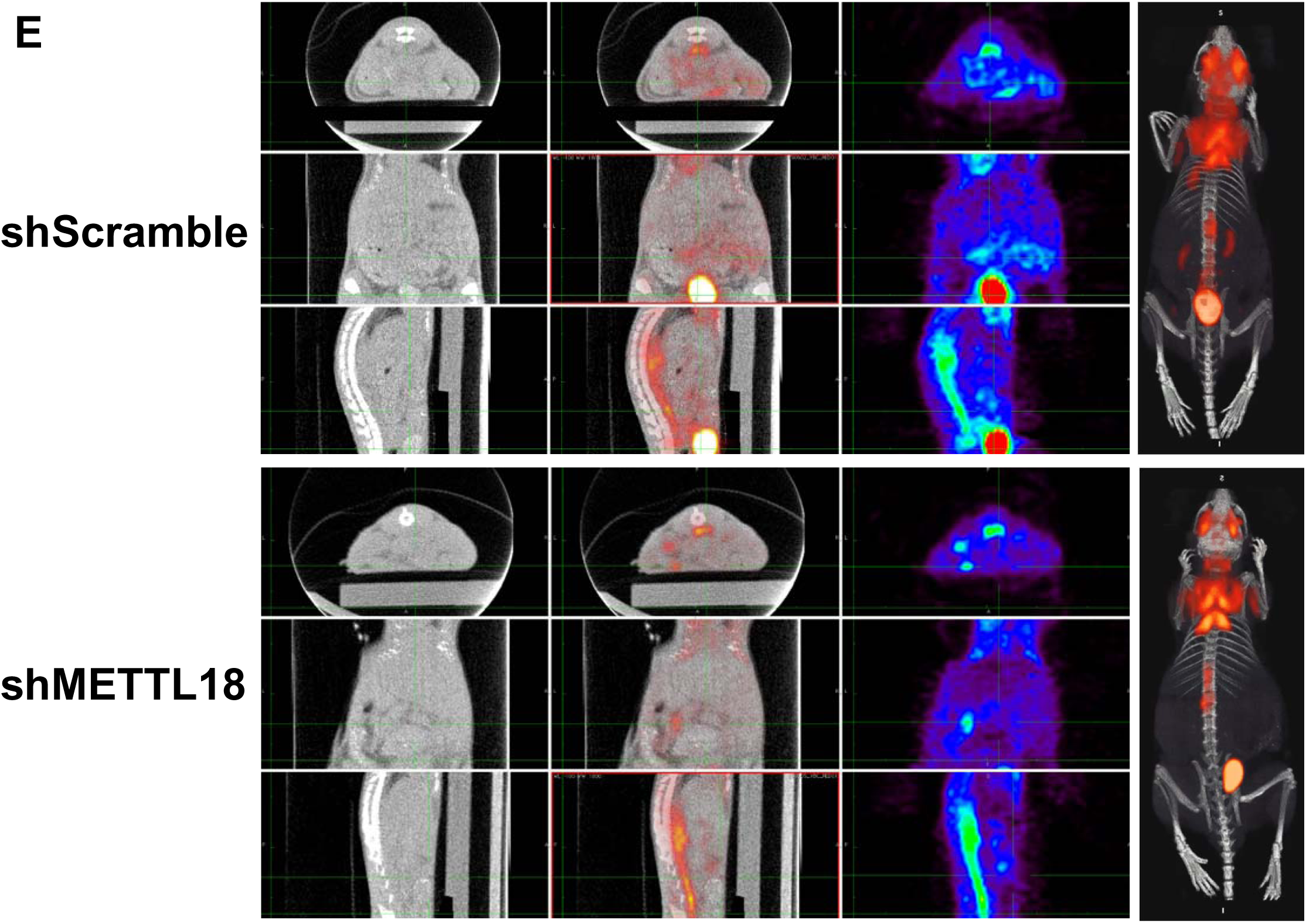
Effect of METTL18 on metastatic responses of breast cancer cells *in vitro* and *in vitro.* (**A**) Protein expression of METTL18 in breast cancer cell lines. Protein levels of HER2, METTL18, and β-actin were identified by Western blot analysis. (**B**) A migration assay was conducted to check the role of METTL18 in cancer cell migration using METTL18 knockdown or METTL18-overexpressing MDA-MB-231 cells. Cell migration range was measured by ImageJ. (**C**) Invasive capacity was investigated in METTL18 knockdown and METTL18-overexpressing MDA-MB-231 cells. Invasion rate was calculated using ImageJ. (**D**) Enzyme activities of MMP-2 and MMP-9 in METTL18 knockdown MDA-MB-231 cells were determined by zymography. (**E**) Metastasis of tumors with intravenously injected METTL18 knockdown MDA-MB-231 cells was visualized via ^18^F-FDG-PET/CT scans.

### Effect of METTL18 on activation of Src kinase and its downstream pathway

To understand the molecular mechanisms by which METTL18 regulates metastatic responses, we examined oncogenic signaling molecules altered by METTL18. Phospho-tyrosine levels of oncogenic proteins, including Syk, JAK2, FGFR1, and c-Raf, were not increased in Myc-METTL18-overexpressed MDA-MB-231 cells (Fig. 3a). In contrast, phosphorylation levels of Src, p85/PI3K, p65/NF-κB, and STAT3 were strongly enhanced in METTL18-overexpressed MDA-MB-231 cells (Fig. 3b Upper left panel). Similarly, phosphorylation of Src, STAT3, and p85 was reduced in shMETTL18-expressing cells (#2975) (Fig. 3b Upper right panel) and in siMETTL18-transfected cells (Fig. 3b Lower left panel) showing clear knockdown levels of METTL18 by its shRNA (Supplementary Fig. S2a Left panel and 2e) and siRNA (Supplementary Fig. S2a Right panel). Consistent with these results, among the various transcriptional factors that play important roles in metastatic responses, only phosphorylation of p65 and p50, downstream transcription factors of Src (Lai et al, 2018), was increased by METTL18 overexpression (Supplementary Fig. S4). In addition, suppressed p-Src, p-p85, and p-STAT3 levels in METTL18-knockout HEK293 cells were recovered with METTL18 transfection in the METTL18-/-HEK293 cells (Fig. 3b, Lower right panel). Expression levels of phosphorylated-Src correlated with METTL18 expression in breast cancer patients from Fig. 1b (Fig. 3c, Supplementary Fig. S5a-c). However, when isoaspartyl, lysine and arginine methyltransferases (PIMT, FAM86A, and PRMT1) were knocked down by their siRNA, p-Src was not decreased (Fig. 3d). Furthermore, survival probability was lower in HER2-negative breast cancer patients with higher Src levels, similar to patients with higher METTL18 level (Fig. 3e). Taken together, these results indicate that the metastasis of breast cancer is regulated by the METTL18-Src signal pathway.

**Fig. 3.**
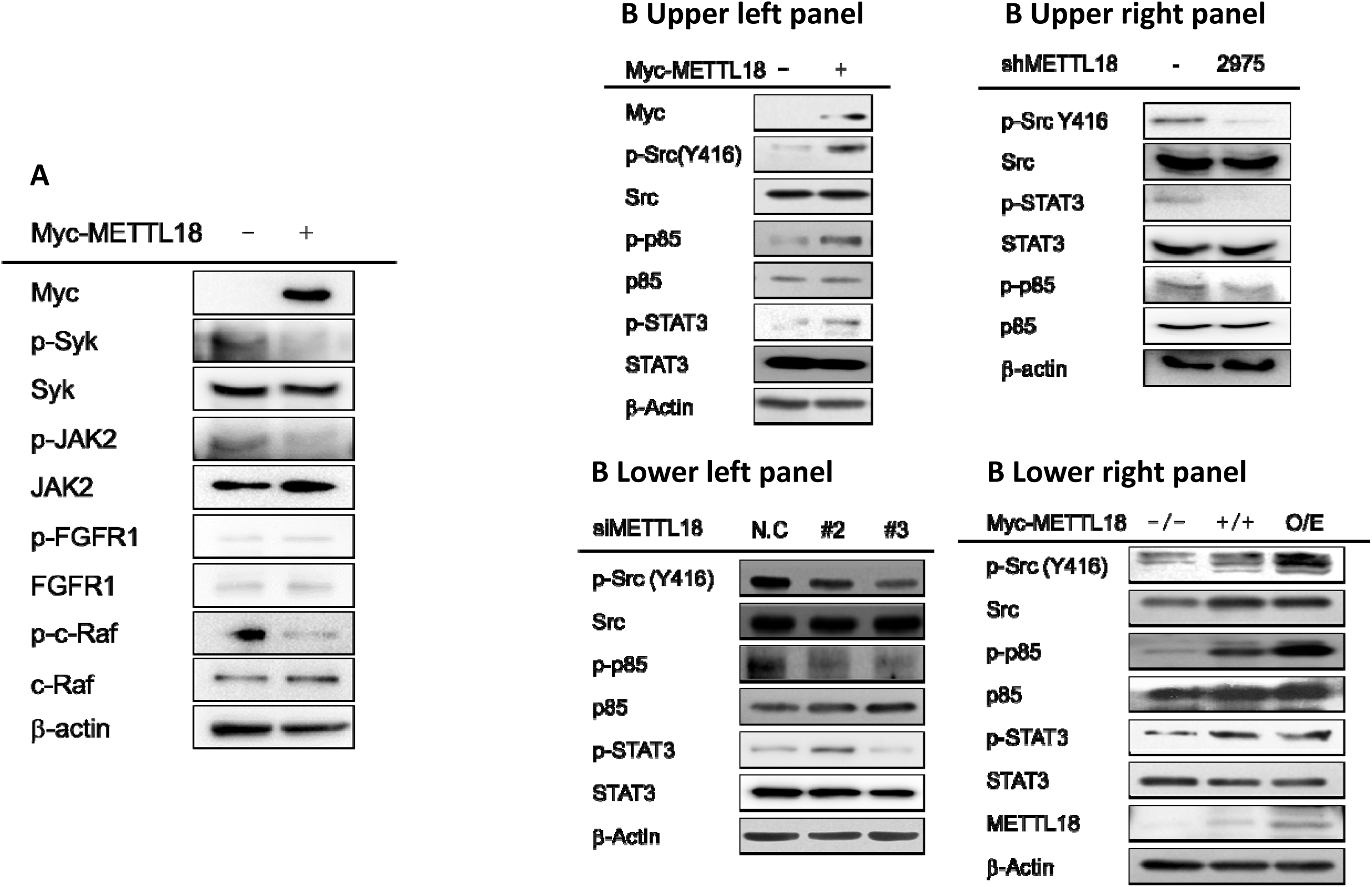
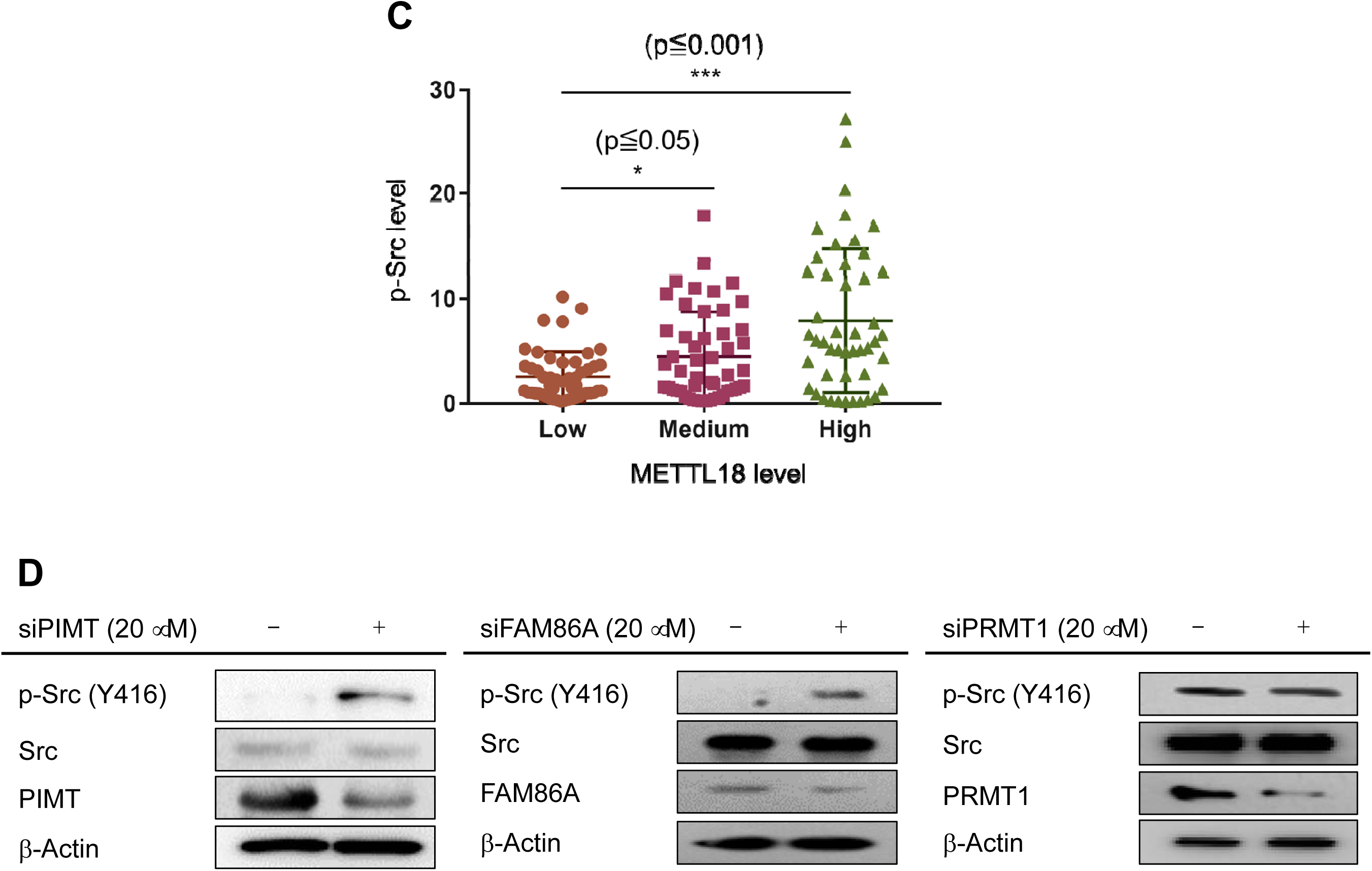
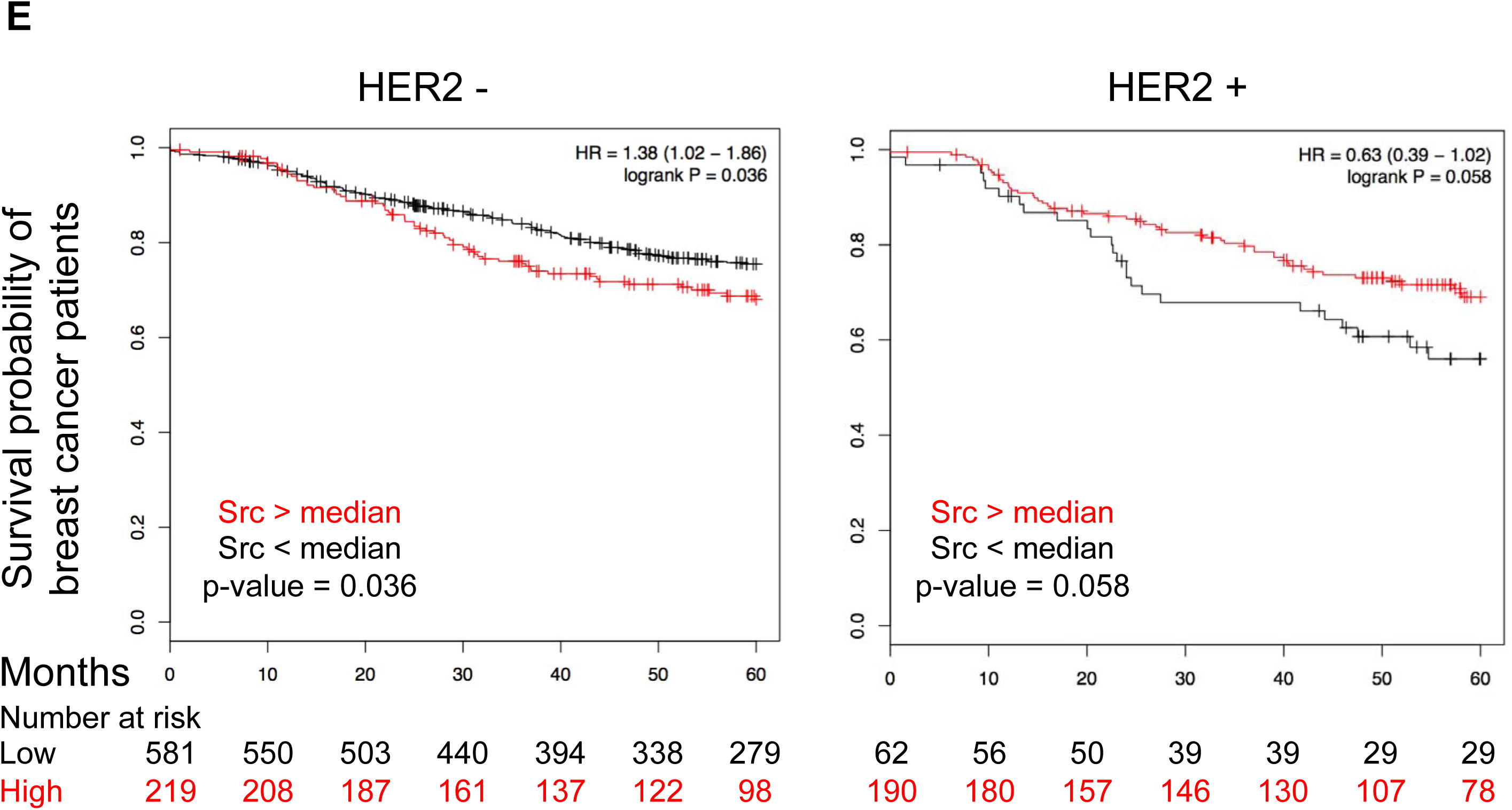
Effect of METTL18 on the activation of Src kinase and its downstream pathway. (**A** and **B**) Total and phosphorylation patterns of tyrosine kinases (Src, Syk, JAK2, and FGFR1), serine/threonine kinases (c-Raf and p85/PI3K), and transcription factors (STAT3) were identified by Western blotting analysis with whole-cell lysates of Myc-METTL18 overexpressing MDA-MB-231 cells (**A** and **B**, Upper left panel), shMETTL18 (**B**, Upper-right panel), siMETTL18 (**B**, Lower-left panel)-expressing MDA-MB-231 cells, and METTL18-knockout HEK cells (**B**, Lower-right panel). (**C**) Correlation between METTL18 and phospho-Src levels was determined from breast cancer patient tissues by Western blotting analysis. (**D**) Western blotting analysis was conducted to detect total and phospho-protein levels of Src in siPIMT-, siFAM86A-, and siPRMT1-transfected MDA-MB-231 cells. (**E**) Kaplan-Meier curve showing survival probability in HER2-negative breast cancer patients with higher or lower Src expression levels (*: p<0.05 and **: p<0.01).

### Involvement of actin in the METTL18-Src regulatory mechanism

To evaluate how METTL18 regulates Src kinase, we investigated the interaction between METTL18 and Src, observing that METTL18 did not bind to Src (Fig. 4a). Moreover, there was no histidine-methylated site in Src according to MALDI-TOF/MS spectrometric analysis (data not shown), indicating that Src is not a direct substrate of METTL18. Therefore, we assessed whether there were molecules capable of regulating the phosphorylation of Src kinase among known binding partners of METTL18, which included RPL3 (de la Cruz et al, 2004), glutamate-rich WD40 repeat-containing 1 (GRWD1) (Clarke, 2018), HSP70 (Cloutier et al, 2013), and actin. Knockdown of RPL3 or GRWD1 or inhibition of HSP70 with VER15508 did not suppress METTL18-induced Src phosphorylation (Fig. 4b). Only siActin reduced the phosphorylation of Src induced by METTL18 to the control level (Fig. 4b), suggesting that actin was involved in METTL18-mediated Src regulation.

**Fig. 4.**
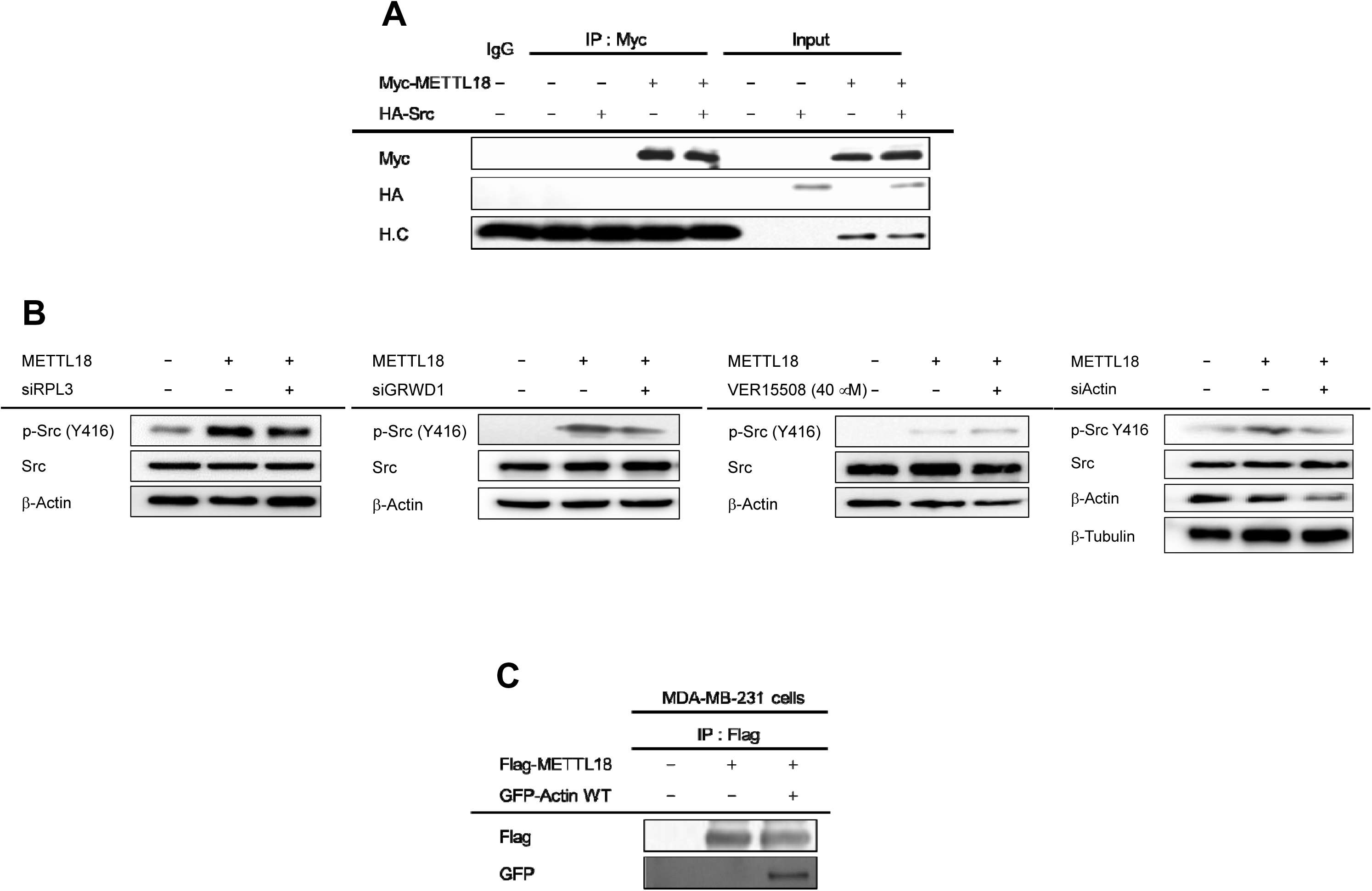
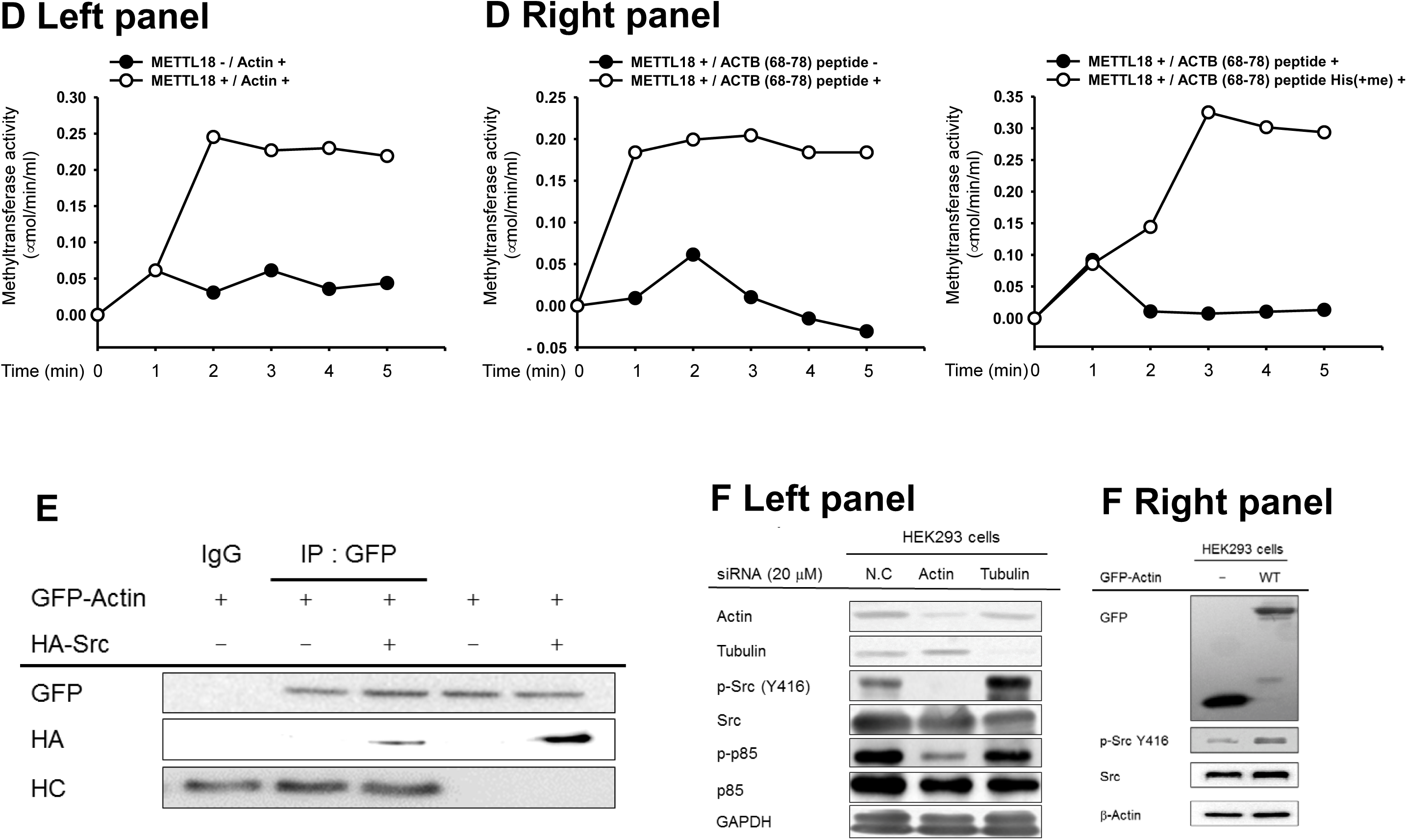
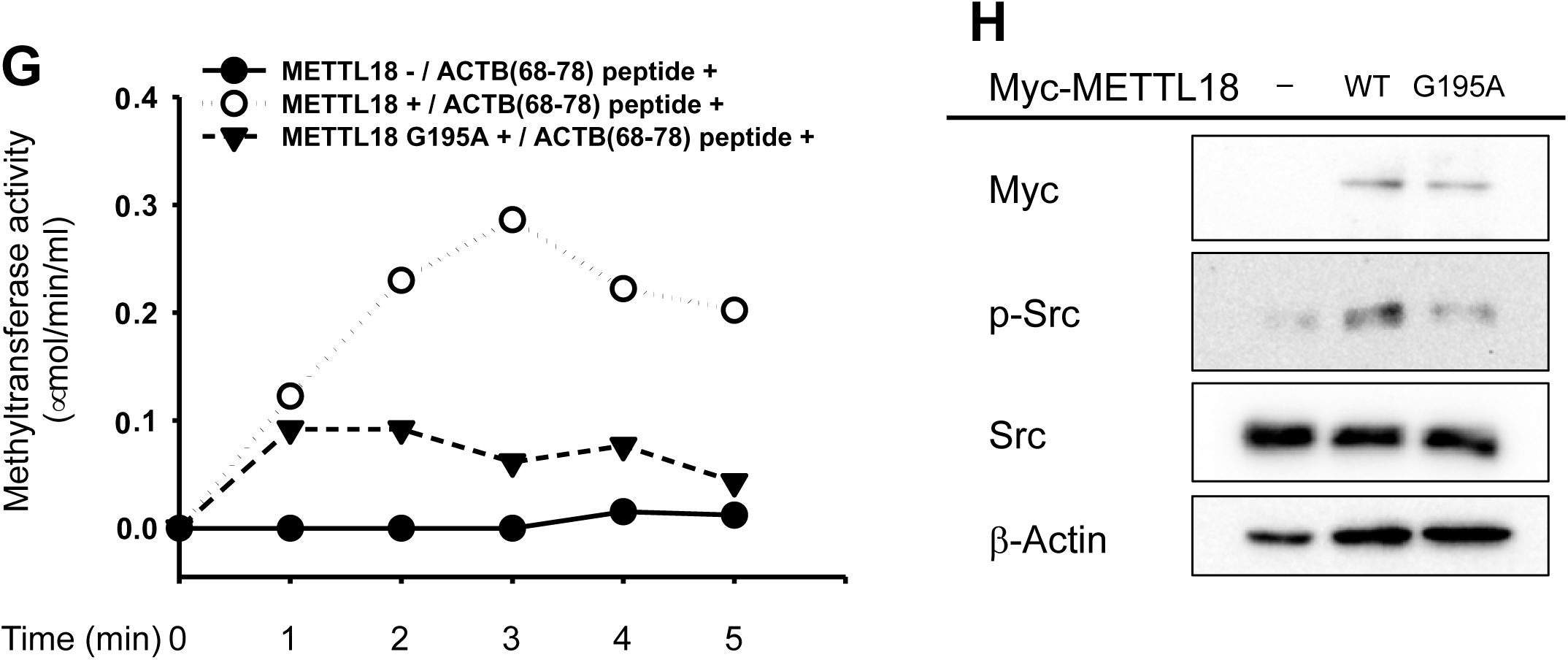
Involvement of actin in the METTL18-Src regulatory mechanism. (**A** and **C**) Interaction levels of METTL18 with Src (**A**) and actin (**C**) were analyzed by immunoprecipitation assay with whole-cell lysates from MDA-MB-231 cells. (**B**) Identification of proteins involved in Src regulation in METTL18 binding proteins was performed by Western blot analysis of METTL18-overexpressing MDA-MB-231 cells during treatment with siRPL3, siGRWD1, VER-15508 and siActin. (**D**) A colorimetric methyltransferase assay was performed using a SAM510:methylation assay kit with immunoprecipitated actin WT or actin-derived histidine-containing peptide. (**E**) Binding of actin with Src was analyzed by immunoprecipitation and Western blotting analysis with whole-cell lysates from MDA-MB-231 cells. (**F**) Total and phospho-form levels of Src, p85/PI3K, tubulin, GAPDH, GFP, and β-actin were determined by Western blotting analysis in HEK293 cells transfected with siActin and siTubulin (**F**, Left panel), or HEK293 cells transfected with GFP-actin-WT (**F**, Right panel). (**G**) A non-radioactive methyltransferase assay was done with immunoprecipitated Myc-METTL18 WT or Myc-METTL18 G195A. ACTB (68-78) peptide was prepared as a substrate source. (**H**) Src and p-Src levels were assessed via Western blotting in MDA-MB-231 cells transfected with Myc-METTL18 WT and Myc-METTL18 G195A.

To provide evidence supporting this hypothesis, we first investigated whether or not METTL18 could methylate actin. Unlike Src (Fig. 4a), actin was able to bind with METTL18 (Fig. 4c). To determine the methyltransferase activity of METTL18 with actin as a substrate source, a non-radioisotope methyltransferase assay was performed. As expected, actin was methylated by METTL18 *in vitro* (Fig. 4d, Left panel). Since the histidine (His) 73 residue is known as a methylation site in actin, 11mer peptides (ACTBs) containing either the His73 residue or methylated His73 were synthesized. As expected, we found that METTL18 methylated the histidine of both actin (Fig. 4d Left panel) and the ACTB (68-78) peptide (Fig. 4d, Right/left panel), whereas the ACTB (68-78) peptide with methylated His73 was not histidine-methylated (Fig. 4d, Right/right panel), indicating that METTL18 could methylate actin at the His73 residue. We then assessed whether actin could regulate Src. Binding analysis indicated that actin bound to Src kinase (Fig. 4e). Depletion of actin but not tubulin in HEK cells inhibited the phosphorylation of Src and p85 (Fig. 4f Left panel), whereas overexpression of actin increased the p-Src level (Fig. 4f Right panel).

Furthermore, to confirm whether METTL18 is a SAM-dependent methyltransferase, we aligned amino acid sequences with previously known protein methyltransferases, including PRMTs 1-10, and identified a conserved sequence (VLDLGCG) within the SAM binding site in METTL18, and a METTL18 and SAM binding structure in the PDB database (Supplementary Fig. S6a, Upper panel, and S6b). Based on this, we constructed METTL18-VLD-AAA and G195A mutants and confirmed that only the mutant containing G195A was expressed properly (Supplementary Fig. S6a Lower panel). Putative binding site of SAM in METTL18 was also speculated by PDB. Thus, G195 was identified as one of amino acids interacting with SAM (Supplementary Fig. S6b). By using CETSA (Martinez Molina et al, 2013) with METTL18-G195A and wild type (Supplementary Fig. S6c), we could confirm that G195 is a critical site for binding to SAM. Overall, actin methylation and p-Src levels were inhibited in METTL18-G196A transfected cells (Fig. 4g and h). These results indicate that METTL18 is methyltransferase to modulate Src kinase through formation of a functional unit with actin.

### Regulatory role of histidine-methylated actin in METTL18/Src-mediated metastatic responses

To analyze the actin-Src regulatory mechanism by METTL18 at the molecular level, we used an actin H73A mutant, since METTL18 methylated the His73 residue of actin (Fig. 4d and Supplementary Fig. S7) (Nyman et al, 2002). Indeed, mass analysis revealed that actin H73A was not methylated, and METTL18 also failed to methylate actin H73A (Fig. 5a and Supplementary Fig. S8a-b). Interestingly, overexpression of actin mutant without H73 residue (Actin-H73A) interfered with actin polymerization in HEK293 cells transfected with GFP-Actin-WT or GFP-Actin-H73A (Fig. 5b) and TLR4-expressing HEK293 cells stimulated with lipopolysaccharide (LPS) (Supplementary Fig. S9), as assessed by confocal microscopy or flowcytometry using florescence dye (phalloidin), which can only bind to polymerized filamentous actin (F-actin) (Lengsfeld et al, 1974). Moreover, the formation of the complex between actin and Src/p-Src/p85 was remarkably failed (Fig. 5c). In addition, when actin polymerization was disrupted by Cyto B treatment, the formation of the actin/Src complex was reduced (Fig. 5d), indicating that the actin/Src complex formation breakdown shown in H73A transfected cells was due to damaged actin polymerization.

**Fig. 5.**
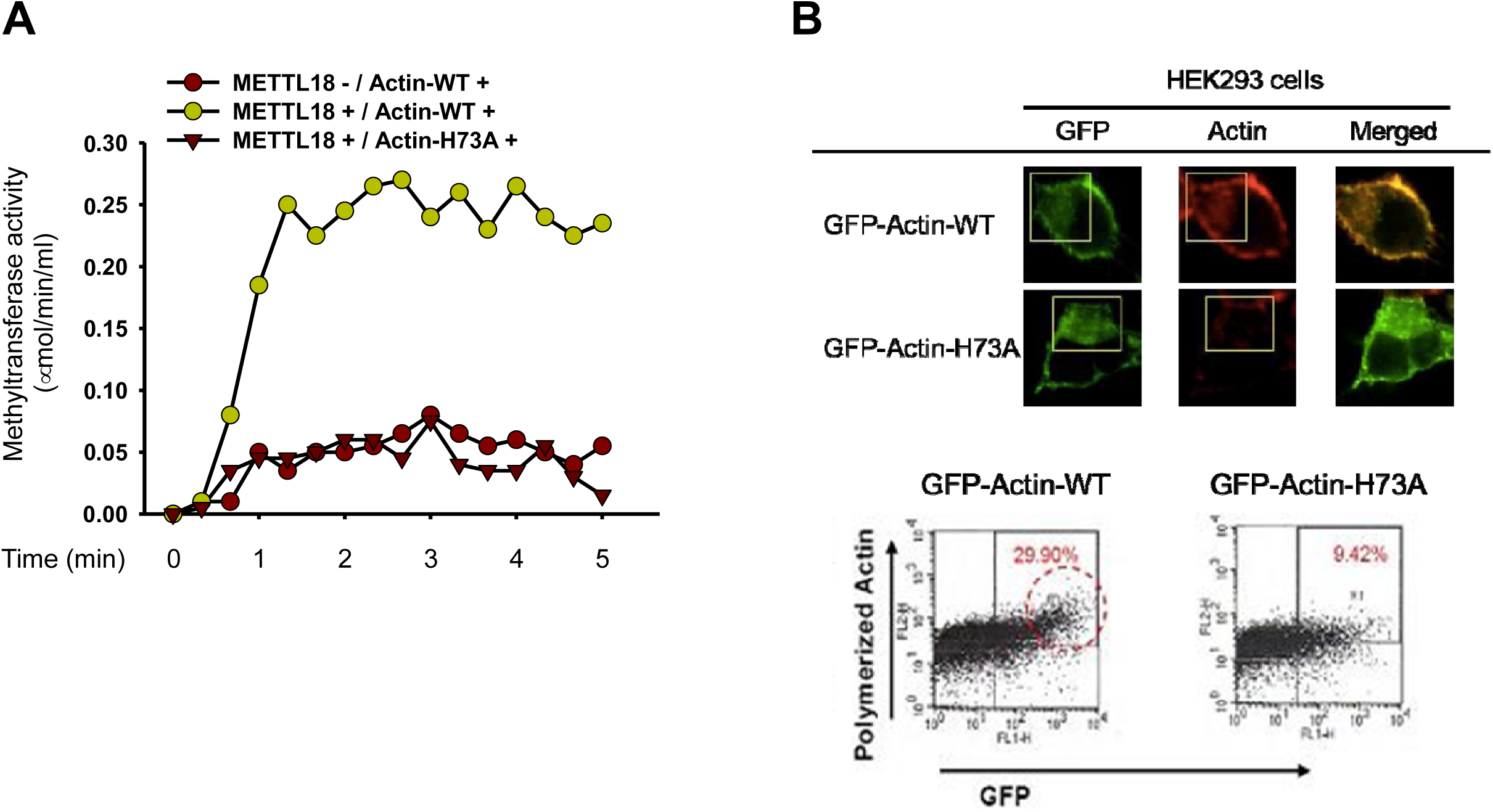
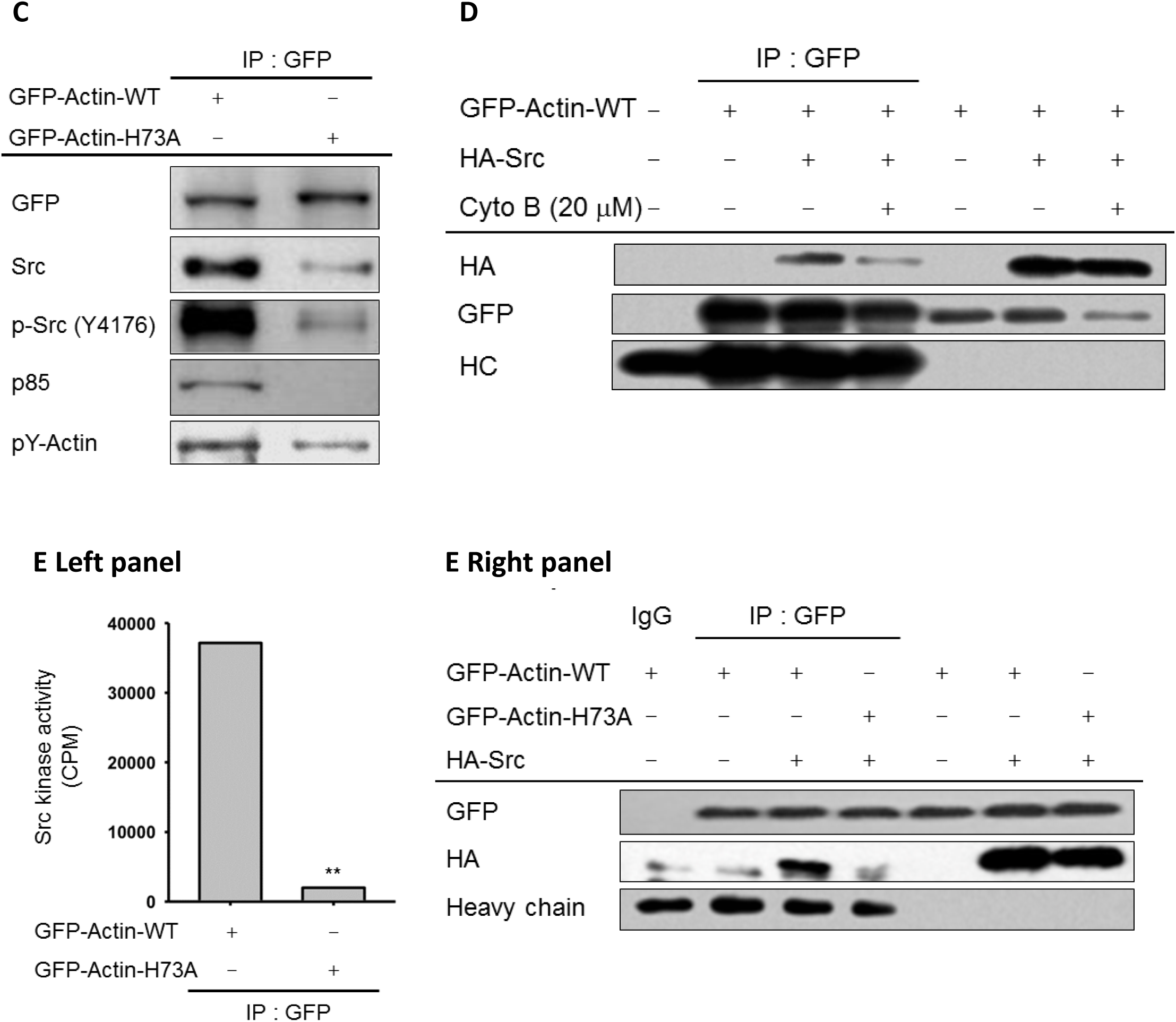
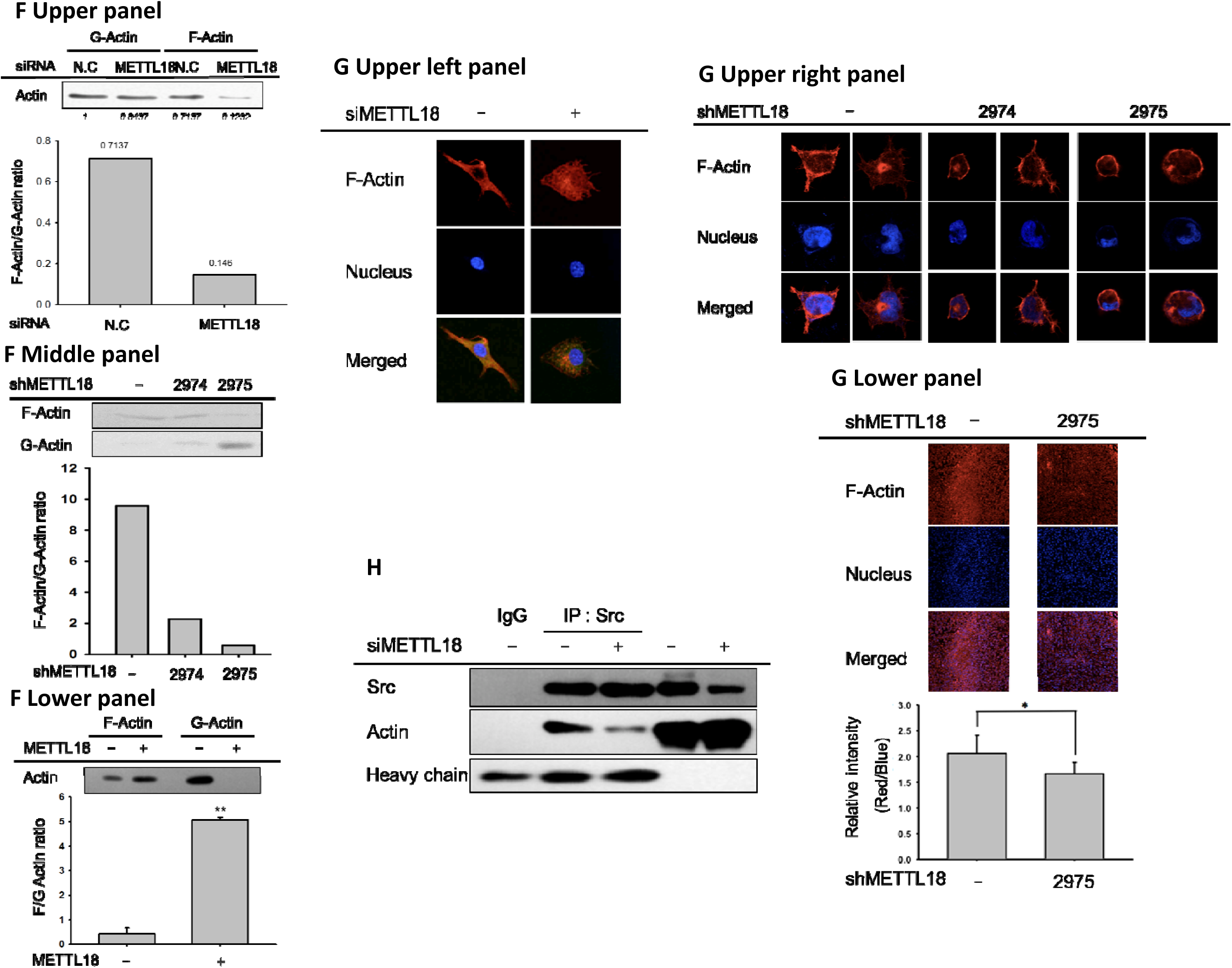
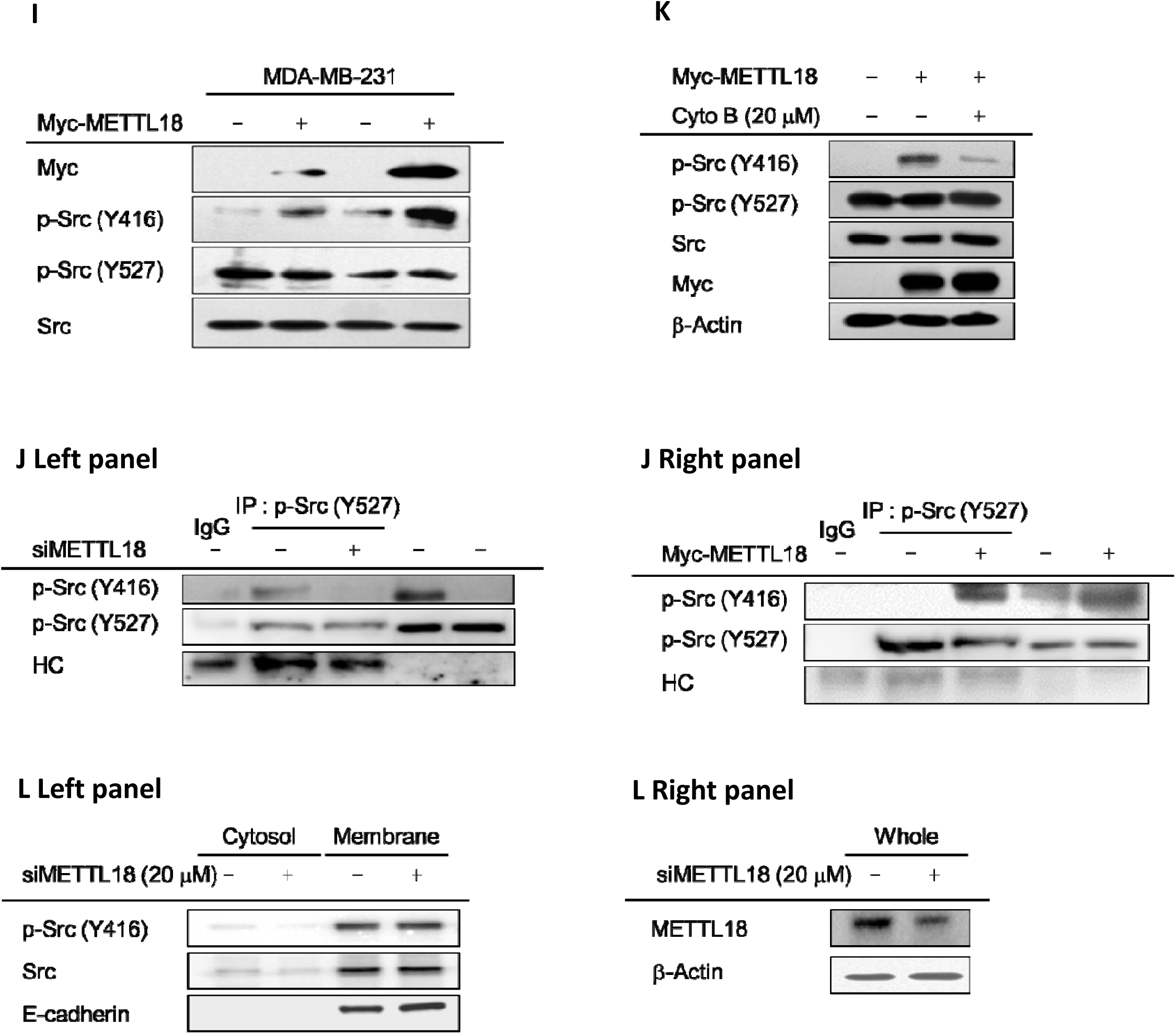
Regulatory role of histidine-methylated actin in METTL18/Src-mediated metastatic responses. (**A**) A colorimetric methyltransferase assay was conducted with immunoprecipitated actin wild type (WT) and actin H73A. An enzyme source was prepared from immunoprecipitated METTL18 in METTL18-overexpressing MDA-MB-231 cell lysate. (**B**) Changes in the actin cytoskeleton were observed by confocal microscopy and flow cytometry in actin WT and actin H73A-transfected cells. Relative intensity was calculated from red and blue fluorescence (Lower panel). (**C** and **E**) Regulatory role of histidine-methylated actin in the activation of Src was examined by immunoprecipitation (**C**) and Src kinase assay (**E**), with whole-cell lysates of MDA-MB-231 cells transfected with actin WT or actin H73A. (**D**) Involvement of actin polymerization in Src/actin complex formation was confirmed by immunoprecipitation with CytoB treatment. (**F**) The role of METTL18 in the stimulation of actin polymerization was determined by an F-actin/G-actin assay in siMETTL18-transfected, shMETTL18-expressing, and METTL18-overexpressing MDA-MB-231 cells. (**G**) Regulatory actions of METTL18 in actin cytoskeleton disruption or polymerization were investigated by confocal microscopy with MDA-MB-231 cells transfected with siMETTL18 (Upper left panel) or shMETTL18 (Upper right panel) and by histological findings with *in vivo* tissue from shMETTL18 cell-injected mice. (**H**) Immunoprecipitation assays were performed in siMETTL18-transfected cells to figure out the function of METTL18 in binding between actin and Src. (**I**) Effects of METTL18 on Src phosphorylation at Y416 and Y527 was determined by Western blotting in MDA-MB-231 cells transfected with Myc-METTL18. (**J** and **K**) Dual phosphorylated form of Src at both Y416 and Y527 residues in MDA-MB-231 cells was detected by immunoprecipitation with anti-p-Src (Y527) antibody and Western blotting analysis with anti-p-Src (Y416) antibody. Alterations in the level of phosphorylated Src at both Y416 and Y527 were observed under siMETTL18, METTL18 overexpression, and CytoB treatment conditions (**K**). (**L**) Localization of phosphorylated Src (Y416) was detected from MDA-MB-231 cells transfected with siMETTL18 by subcellular (cytosol and membrane) fractionation and Western blotting analysis.

Because phosphorylation of actin at tyrosine residues was also decreased in the cells with actin H73A (Fig. 5c), we examined whether other post-translational modifications of actin affected complex formation with Src. However, the phosphorylation, glutathionylation, and ubiquitinylation of actin were not involved in actin/Src binding (Supplementary Fig. S10). Furthermore, lower enzyme activity of Src kinase as well as its biding level to GFP-H73A were observed (Fig. 5e), and reduced levels of metastatic responses, including migration and invasion, were also seen in the H73A transfection cells (Supplementary Fig. S11a-b). Similar to the H73A transfection results, F-actin reduction, actin rearrangement disruption, and morphological changes were observed with METTL18 depletion by shMETTL18 (#2974 and 2975) and siMETTL18 (#3) (Fig. 5f, Upper and middle panels, and 5g, Upper panel). Levels of polymerized actin stained with phalloidin were also found to be decreased in tumor tissues expressing shMETTL18 *in vivo* (Fig. 5g, Lower panel). Conversely, F-actin was increased when METTL18 was overexpressed (Fig. 5f, Lower panel). The interaction between actin and Src was suppressed in METTL18 knockdown cells transfected with siMETTL18 (#3) (Fig. 5h), but binding with CSK and PTP1B was not influenced (Supplementary Fig. S12). These results imply that METTL18 induces the methylation and actin polymerization of the actin His73 residue, resulting in the formation of a functional unit containing Src/p85, centered on actin.

For an in-depth understanding of the Src regulatory mechanism by METTL18, we compared the two forms of Src kinase, inactive and active. Generally, phosphorylated Src at tyrosine 416 is referred to as the active form and p-Src (Y527) is the inactive form (Roskoski, 2005). Intriguingly, phosphorylation of Src at Y416 was induced by METTL18 overexpression while p-Src (Y257) did not change (Fig. 5i). These results suggested the possibility that a new form of Src kinase might be present in addition to the inactive and active forms, and, in fact, we confirmed the presence of Src kinase phosphorylated at both sites by immunoprecipitation with antibody to p-Src (Y527) and Western blotting with antibody to p-Src (Y416) (Fig. 5j, Left panel). Moreover, overexpressed METTL18 introduced a new phosphate group into the tyrosine 416 residue of Src kinase in which tyrosine 527 was phosphorylated (Fig. 5j, Right panel). Treatment with Cyto B also blocked phosphorylation of Y416 independent of Y527 phosphorylation (Fig. 5k), suggesting that actin polymerization participates in the modulation of Src phosphorylation. Moreover, siMETTL18 (#3) blocked the level of cytosolic phospho-Src at Y416 but not membrane Src (Fig. 5l), implying that only cytosolic Src is modulated by the METTL18/actin unit.

### Role of HSP90AA1 in METTL18-mediated actin polymerization and Src activation

In addition, based on the previous finding that METTL18 can associate with HSP90 and is involved in regulating actin dynamics (Taiyab & Rao Ch, 2011), we also tested whether HSP90 is part of this functional unit to regulate the actin cytoskeleton. Interestingly, overexpression of HSP90AA1 increased the binding level between METTL18 and actin (Supplementary Fig. S13a), whereas knockdown of HSP90AA1 strongly decreased their binding (Supplementary Fig. S13b-c). In addition, like Cyto B and siMETTL18, siHSP90AA1 or chemical inhibitor of HSP90AA1 (17-AAG) but not HSP70 inhibitor (VER-15508) reduced METTL18-mediated Src phosphorylation, actin polymerization, and cellular tumorigenic responses such as cell migration, invasion, and morphological changes in MDA-MB-231 cells overexpressed with Myc-METTL18 or empty vector (Fig. S13d-i).

### Comparison of Src activation property of actin histidine methyltransferases, METTL18 and SETD3

Since SETD3 has also been reported as an enzyme that methylates actin at H73 (Wilkinson et al, 2019), we examined whether SETD3 could regulate Src activity. Interestingly, SETD3 did not affect p-Src level, although it induced actin methylation at a level similar to METTL18 (Supplementary Fig. S14a-b). To discriminate such difference between METTL18 and SETD3, whether SETD3 can bind to HSP90AA1, a key molecule to control actin/Src interaction (Supplementary Fig. S13), was examined. Interestingly, unlike METTL18 (Supplementary Fig. S13a), there was no association between SETD3 and HSP90AA1, according to immunoprecipitation and Western blotting analysis (Supplementary Fig. S14c). In addition, the distribution patterns in ER and induction of p-Src level under overexpression of Myc-METTL18 or Flag-SETD3 were found to be distinct between METTL18 and SETD3 by confocal microscopy (Supplementary Fig. S14d-e). These results suggest that METTL18 modulates actin in a different way than SETD3, resulting in a different ability to control Src kinase. Interestingly, the mRNA expression pattern of METTL18 and SETD3 in breast cancer patients showed the opposite (Supplementary Fig. S14f), indicating that METTL18 could be more pathophysiological in tumorigenesis of breast cancer.

## Discussion

About 50 years ago, Perry and colleagues showed that histidine residues in proteins could be methylated in mammalian, fish, and bird skeletal muscles (Johnson et al, 1967). Thus far, researchers have focused more on lysine and arginine methylation than on histidine methylation (Bedford & Richard, 2005; Kubicek & Jenuwein, 2004). However, a recent study identified a rat actin-specific histidine N-methyltransferase known as SETD3, which is a protein-histidine methyltransferase that has been found in vertebrates and the first SET-domain-containing enzyme exhibiting dual methyltransferase specificity toward both Lys and His residues in proteins. SETD3 is also responsible for the methylation of H73 in human and Drosophila actin (Kwiatkowski et al, 2018). Here, we have investigated the biological role of METTL18, another protein histidine methyltransferase, in association with cancer metastasis. Despite the higher levels of METTL18 in other cancers, including bladder, liver, and lung adenocarcinomas, studies have mainly focused on breast cancer because the expression of METTL18 is associated with survival rate in HER2-negative breast cancer patients (Fig. 1b, 1c, and 1d). In the absence of METTL18 generated using shMETTL18, the metastasis properties of HER2-negative MDA-MB-231 cells were reduced (Fig. 2). Activity of both MMP-2 and MMP-9 was also diminished under METTL18 knockdown conditions (Fig. 2d). Moreover, cancer metastasis was diminished in shMETTL18-expressing MDA-MB-231 cell-injected mice *in vivo* (Fig. 2e). Together, these *in vitro* and *in vivo* experiments demonstrate that METTL18 plays a critical role in the metastasis of HER2 negative breast cancer.

Src kinase is a well-known oncogenic protein, and a critical role for Src activation in promoting brain metastasis of breast cancer cells has been well-defined (Fan et al, 2019; Lamar et al, 2019). Indeed, blockade of Src gave us a promising activities to suppress metastatic potential of breast tumor cells. It has been also reported that Src kinase-dominant negative (DN) reduces cell proliferation and migration in breast cancer (Gonzalez et al, 2006). In this study, it was revealed that METTL18 regulates the phospho-level of Src, as well as STAT3 and p85, which are downstream molecules of Src (Fig. 3a and 3b). Furthermore, since Src kinase is involved in MMP activity (Mon et al, 2017), we conclude that the induction of metastatic responses by METTL18 is due to the alteration of Src kinase. In support of our argument, we found that increased p-Src level was directly proportional to METTL18 expression in the tissues of breast cancer patients (Supplementary Fig. S1a). Methyltransferases such as PIMT, FAM86A, and PRMT1 did not influence the phosphorylation of Src, suggesting that METTL18 has specificity in regulating Src kinase (Fig. 3d).

Since Src does not bind to METTL18 (Fig. 4a) and also lacks a methylated site (data not shown), we sought to find molecules capable of regulating Src kinase among the direct binding partners of METTL18. Actin was found to be involved in METTL18-mediated Src activation (Fig. 4b) among HSP70, GRWD1, and RPL3 (Cloutier et al, 2013). In addition, the inhibition of Src kinase activity has been reported in breast cancer cells when actin polymerization is disturbed (Kim et al, 2013). Therefore, we hypothesized that METTL18 could regulate Src kinase through actin. We observed that Src kinase activity induced by METTL18 was reduced to almost basal levels after siActin transfection (Fig. 4f Left panel). In addition, METTL18 methylated His73 in actin (Fig. 4d and Supplementary Fig. S8a and b). The most prominent outcome of actin methylation is actin polymerization (Nyman et al, 2002). Therefore, we attempted to understand the mechanism underlying regulation of Src by METTL18 by examining the relationship between actin polymerization and Src kinase.

To date, the role of actin polymerization in eukaryotic cells has been limited to cell shape determination, movement, division, secretion, and providing pathways for pathogen invasion (Carpenter, 2000). Although studies have been conducted on signal transduction pathways that regulate actin polymerization and contraction (Carpenter, 2000), those on signaling pathways that are controlled by actin polymerization are rare. According to this study, signal transduction could be induced by conjugating signaling molecules to polymerized actin. In particular, F-actin is thought to contribute to complex formation with Src kinase/PI3K (Fig. 4b, 4f Left panel, and 5c). As siMETTL18 transfection did not inhibit the binding ability of PTP1B (Bjorge et al, 2000) and CSK (Okada, 2012) to actin (Supplementary Fig. S12), it is believed that F-actin as well as HSP90AA1 provide a scaffolding structure for binding between Src and Src substrates (Supplementary Fig. S12 and S13). These results indicate that METTL18 forms a functional complex for actin regulation and, ultimately, that METTL18 regulates Src kinase through actin methylation. Interestingly, SETD3, a known actin histidine methyltransferase (Wilkinson et al, 2019), was not involved in Src regulation (Supplementary Fig. S14). In a comparative analysis of actin regulatory mechanisms, we observed that SETD3, but not METTL18, was unable to induce Src phosphorylation (Supplementary Fig. S14a), although both SETD3 and METTL18 increased histidine-methylation in ACTB peptide (Supplementary Fig. S14b). Interestingly, the mRNA expression pattern of METTL18 and SETD3 in breast cancer patients and their p-Src activation level in MDA-MB-231 cells showed the opposite (Supplementary Fig. S14f and 14e). Further, SETD3 did not bind to HSP90AA1 (Supplementary Fig. S14c). These results suggested that METTL18, in cooperation with a complex composed of actin polymerization, HSP90AA1, and Src, might have more pathophysiological function than SETD3.

Interestingly, METTL18 induced phosphorylation of Y416 without Y527 dephosphorylation of Src kinase; in this process, Src kinase with Y416 and Y527 both phosphorylated was found (Fig. 5i, 5j, and 5k). As the presence of this type of Src kinase cannot be explained by the conventional Src regulatory mechanism, it is presumed that Src regulation by METTL18 would follow a novel pathway. Recently, an intermediate form of Src was proposed through Src structural analysis using a computational paradigm combined with transition pathway techniques and a Markov state model (Shukla et al, 2014). According to their findings, tyrosine 416 is phosphorylated in the closed repressed conformation of c-Src, which suggests a model in which the tyrosine 416 of c-Src is phosphorylated even though tyrosine 527 is also phosphorylated. Src kinase that is closed (or phosphorylated at Y527) and phosphorylated at Y416 is defined as an intermediate form (Shukla et al, 2014). Moreover, phosphorylated Y527 stabilizes a closed conformation and Y416 phosphorylation in the closed conformation (p-Y527) could occur due to auto-phosphorylation, depending on the self-association of the Src kinase itself (Camara-Artigas et al, 2014). Accordingly, we propose a novel Src regulatory model, in which the Src conformation is changed from the closed intermediate form to the open form where Src kinase associates with methylated actin, after which Src is activated (Fig. 6). Furthermore, in. our experiments, siMETTL18 blocked the levels of cytosolic phospho-Src at Y416, but not membrane Src (Fig. 5l Left panel), suggesting that only cytosolic Src is modulated by the METTL18/actin unit.

**Fig. 6.**
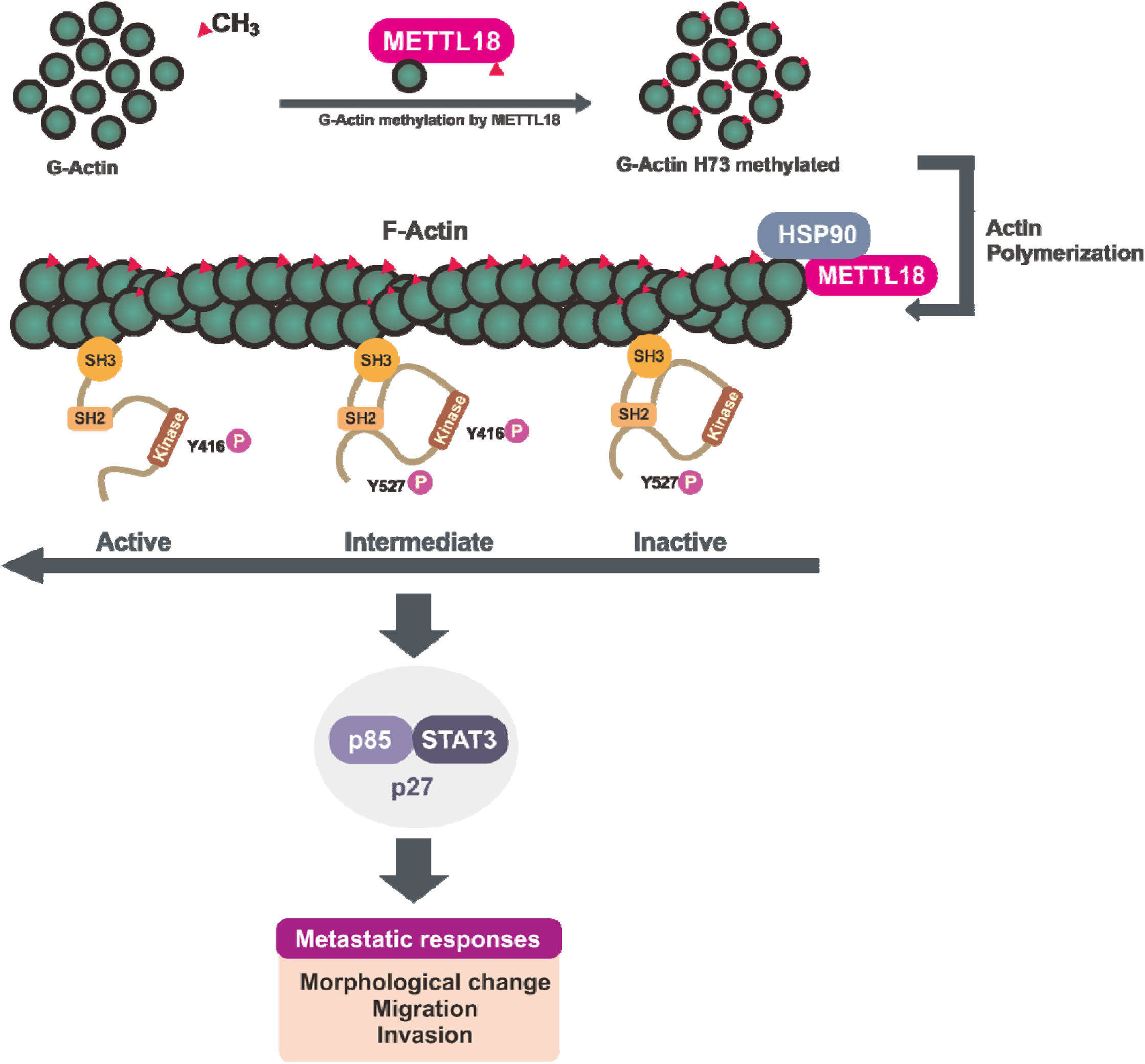
Schematic diagram of METTL18-mediated metastatic responses in breast cancer.

In summary, METTL18 was highly expressed in HER2-negative breast cancer, and breast cancer patients with increased levels of METTL18 have a poor prognosis. In addition, METTL18 has been implicated in the metastasis of breast cancer both *in vivo* and *in vitro*. Moreover, as summarized in Figure 6, we demonstrated that METTL18 regulates the methylation and polymerization of actin through a complex formation with HSP90AA1, eventually regulating Src kinase activity. These results are expected to provide therapeutic targets for HER2-negative breast cancer, especially in treatment of TNBC.

## Materials and methods

### Materials, construction of expression vectors and virus production

Detailed information regarding the antibodies, construction of expression vectors, virus production and other related materials is described in Supportive information.

### Cell culture

HEK293 and MDA-MB-231 cells purchased from ATCC (Manassas, VA, USA) were maintained in Dulbecco’s modified Eagle’s medium (DMEM; Hyclone, Logan, UT, USA) supplemented with 10% FBS and penicillin/streptomycin (P/S) at 37°C in a humidified atmosphere containing 5% CO_2_. Mycoplasma contamination in cells was regularly checked with the BioMycoX Mycoplasma PCR Detection Kit (CellSafe, Seoul, Korea). Stable cells expressing shMETTL18s were infected by lentiviral particles in the presence of polybrene (8 μg/ml). Treatment with puromycin (1 μg/ml) for two days after infection was used to select knockdown or knockout cells. The expression level of METTL18 was confirmed by real-time PCR.

### Human breast tumor tissue preparation

Human breast tumor tissue and patient overall survival were obtained from Samsung Medical Center (Seoul, Korea). This study was approved by the Institutional Review Board at Sunkyunkwan University Medical School. Whole lysates of tumor tissues were used to detect METTL18, p-Src, Src, and β-actin levels by immunoblotting.

### Protein isolation and Western blotting

For lysis, cultured cells (5 × 10^6^ cells/ml of HEK293 and MDA-MB-231 cells) or tissues (10 mg of normal or cancer tissues) were treated with lysis buffer. To prepare membrane fractions, cell lysates were centrifuged at 19,326 × g for 1 min. The pellets contained nuclear material, and so the supernatant was further centrifuged at 14,000 rpm for 1 h at 4°C to make membrane and cytosolic fractions as the second step. Finally, the pellet was also treated with extraction buffer (without Triton X-100). The levels of proteins from whole lysates, membrane fractions or nuclear extracts were analyzed by Western blotting through separating proteins on 10% or 12% SDS-polyacrylamide gels, transferring the proteins to polyvinylidenedifluoride (PVDF) membranes, blocking the membrane in Tris-buffered saline containing 3% bovine serum albumin, and detecting with ECL solution (Amersham, Little Chalfont, Buckinghamshire, UK).

### Immunoprecipitation

Cell lysates with equal amounts of protein (500 μg for exogenous protein, 1000 μg for endogenous protein) were incubated with 5 μl of anti-Myc, Flag, GFP, Actin, Src, HA, or GFP antibodies overnight at 4°C. The immune complexes were mixed with 20 μl of Protein A or G sepharose 4 Fast flow beads (50% v/v, GE Healthcare Life Sciences, Marlborough, MA, USA). Whole cell lysates and immunoprecipitates were detected by Western blotting. Mouse TrueBlot® ULTRA: Anti-Mouse Ig HRP and Rabbit TrueBlot®: Anti-Rabbit IgG HRP (Rockland Immunochemicals, Pottstown, PA, USA) was used to detect to prevent overlap with IgG-heavy chain.

### Reverse transcription and real-time polymerase chain reaction (PCR)

Total RNA was extracted with TRIZOL (GIBCO-BRL, Grand Island, NY, USA). After preparing cDNAs with MMLV RTase (SuperBio, Daejeon, Korea), semiquantitative polymerase chain reaction (PCR) and real-time PCR were performed. The semiquantitative and quantitative PCR primers are listed in Supporting Materials and Methods (Supplementary Table 1). All primers were purchased from Macrogen, Inc. (Seoul, Korea).

### Determination of metastatic potential *in vitro*

Morphological changes were observed by taking photos with confocal and inverted microscopes. For the wound healing (cell migration) assay, a wound was generated with monolayer-grown MDA-MB-231 cells by scraping with a p200 pipette tip. For the invasion assay, the invasive capacity of MDA-MB-231 cells was determined using matrigel-coated plates. The cells were then fixed in 4% formaldehyde; subsequently, hematoxylin and eosin staining were used to count the number of cells that successfully penetrated the matrigel layer. A clonogenic assay was carried out with 1,000 cells in 6-well plates. Cells were cultured for 14 days and then stained with crystal violet.

### Gelatin zymography

To measure the activity of MMP-9 and MMP-2 in the conditioned medium, zymography was employed. For this experiment, a sample mixture of 10 μg protein and non-reducing sample buffer was loaded on a polyacrylamide gel containing gelatin (0.25 mg/ml; Sigma). The gel was then soaked (36 h, 37°C) in a renaturing buffer. Coomassie brilliant blue and methanol/acetic acid (30%/10% v/v) solution were used for staining and destaining of the gel.

The lower bands were gelatinase-A (MMP-2) at 60 kD and 72 kD, while the upper bands were gelatinase-B (MMP-9) at 95 kD.

### Tumor xenografts, metastasis assays, and PET/CT

Animals were cared for following the Guide for the Care and Use of Laboratory Animals and mouse studies were carried out following procedures approved by the Institutional Animal Care and Use Committee at the National Cancer Center (Ilsan, Korea). For inoculation into nude mice, 5×10^6^ indicated cells were injected subcutaneously on the dorsal surface of 6-week-old female athymic *nu/nu* mice (Harlan Sprague-Dawley, Indianapolis, IN, USA) for the xenograft study and intravenously in the tail vein for the tumor metastasis assay. Tumor volumes and body weights were monitored every other day. For the tumor metastasis assay, tumor metastasis was detected by PET/CT scanning. PET and CT images are presented and regenerated by MMWKS Vista-CT software provided through the scanner manufacturer.

### G-actin/F-actin assay

The G-actin/F-actin assay was performed with a G-actin/F-actin *in vivo* assay kit (Cytoskeleton, Inc., Denver, CO, USA) according to the manufacturer’s instructions. To evaluate the ratio of G-actin to F-actin, the same volumes of each fraction were loaded onto a 10% SDS-PAGE and analyzed by Western blotting with anti-actin antibody.

### Confocal microscopy

For confocal microscopy, shMETTL18-overexpressing, siMETTL18-48 h-transfected or GFP-actin-WT and -H73A-48 h-transfected MDA-MB-231 cells were prepared before plating. A density of 2 × 10^5^ cells were seeded in 12-well plates over sterile cover slips. The cells were washed twice with 1 ml PBS and fixed with 3.7% paraformaldehyde. After washing three times with PBS, the coverslips were blocked in 1% BSA. For staining the cells, rhodamine phalloidin (Invitrogen, 1:40), Hoechst (1:1000) and indicated antibodies were used to stain polymerized actin, nucleus, and specific target proteins. Coverslips were mounted onto slides using fluorescent mounting medium (DakoCytomation, Carpentaria, CA, USA). The cells were viewed by confocal microscopy (LSM800, Zeiss).

### Mass spectrometry

To identify actin histidine methylation sites, 1.5 x 10^6^ cells were plated in a 6 cm plate and transfected with GFP-actin-WT or GFP-actin-H73A. The cells were prepared by immunoprecipitation with anti-GFP antibody. Immunoprecipitates were separated by protein size using SDS-PAGE, and the gels were stained with Coomassie blue. Bands of interest (∼70 kDa) were cut from the gels and analyzed by mass spectrometry.

### Methyltransferase activity assay and Src family kinase assay

To evaluate methyltransferase activity, immunoprecipitated METTL18 was used as an enzyme source, and immunoprecipitated Actin-WT or Actin-H73A, or histidine-containing peptide derived from actin were utilized as substrate sources. A non-radioactive colorimetric continuous enzyme kit (#786-430; G-Biosciences, St. Louis, MO, USA) was used to measure the enzyme activity of METTL18 according to the manufacturer’s instructions. Src kinase activity was measured with immunoprecipitated Src in reaction buffer using a Src kinase assay kit (Upstate Biotechnology, Inc., Lake Placid, NY, USA) in accordance with the manufacturer’s protocol. A substrate peptide (KVEKIGEGTYGVVYK) designed from p34cdc2 was used.

### Kaplan-Meier analysis

Breast Cancer Gene-Expression Miner v3.0 software designed by *Jezequel et al.* (Jézéquel et al, 2012) was used for Kaplan-Meier analysis with respect to HER2 status.

### Statistics and reproducibility

Unless otherwise stated, all data are presented as the mean ± S.E.M. of at least three independent experiments. Differences were analyzed by Kruskal-Wallis/Mann-Whitney test. All P values ≤ 0.05 were considered statistically significant. GraphPad Prism version 6.0 (GraphPad Software) and SPSS (SPSS Inc., Chicago, IL, USA) were used for data analysis.

## Abbreviations

ACTB: Actin, cytoplasmic 1
AdOx: Adenosine dialdehyde
AsTPs: Arsenic-transactivated protein 2
CytoB: Cytochalasin B
ER: Estrogen receptor
FGFR1: Fibroblast growth factor receptor 1
GFWD1: Glutamate-rich WD40 repeat-containing 1
HER2: Human epidermal receptor 2
His: Histidine
HSP90: Heat shock protein 90
JAK2: Tyrosine-protein kinase JAK2
METTL18: methyltransferase-like protein 18
PCR: Polymerase chain reaction
PIMT: Protein L-isoaspartyl methyltransferase
PKMT: Protein lysine methyltransferase
PR: Progesterone receptor
PRMT: Protein arginine methyltransferase
RPL3: 60S ribosomal protein L3
Src: Proto-oncogene tyrosine-protein kinase Src
STAT3: Signal transducer and activator of transcription 3
Syk: Tyrosine-protein kinase SYK
TNBC: Three negative breast cancer
VEGFA: Vascular endothelial growth factor A

## Author contributions

H.G.K, J.H.K., and J.Y.C. conceived and designed the experiments. H.G.K, J.H.K. W.S.Y., J.G.P., Y.G.L., E.K., S.H.K., N.Y.S., Y.S.Y., M.K.J., C.Y.L., Z.R.A., B.C.Y., and S.U.K. performed the experiments. H.G.K, J.H.K. W.S.Y., Y.B.K., J.W.H., and J.Y.C. supervised data collection, performed the analysis, and wrote the manuscript. All authors discussed the results and commented on the manuscript.

## Funding

This research was supported by the Basic Science Research Program through the National Research Foundation of Korea (NRF) funded by the Ministry of Education (Grant No.: 2011-0016397 and 2017R1A6A1A03015642 to J.Y.C.), Republic of Korea.

## Availability of data and materials

The data used or analyzed during this study are included in this article and available from the corresponding author upon reasonable request.

## Ethics approval and consent to participate

All procedures performed in studies involving human participants were in accordance with the ethical standards of the Research Ethics Committee of Samsung Hospital and Sungkyunkwan University and with the 1964 Helsinki declaration and its later amendments. Written informed consent to participate in the study and for sample collection was obtained from breast cancer patients.

## Consent for publication

All subjects signed written informed consent.

## Competing interests

The authors have no competing interests to declare.

## Supportive information

### Materials and Methods

#### Materials and Antibodies

Anti-METTL18 (Atlas HPA035314), anti-HER2 (CST 2242), anti-β-actin (CST 4967), anti-phospho-Tyrosine (Upstate 05-321), anti-Myc (CST 2276), anti-phospho-Src-Y416 (CST 2101), anti-phospho-Src-Y527 (CST 2105), anti-Src (CST 2109), anti-phospho p85 (CST 4228), anti-p85 (CST 4292), anti-phospho-Syk (CST 2711), anti-Syk (CST 2712), anti-phospho JAK2 (CST 3771), anti-JAK2 (CST 3230), anti-phospho-FGFR1 (CST 3471), anti-FGFR1 (CST 3472), anti-phospho-c-Raf (CST 9427), anti-c-Raf (CST 9422), anti-phospho-p65 (CST 3039), anti-p65 (CST 8242), anti-phospho-p50 (SC 33022), anti-p50 (CST 12540), anti-phospho-c-Jun (CST 9164), anti-c-Jun (CST 9165), anti-phospho-c-Fos (CST 5348), anti-c-Fos (CST 2250), anti-phospho-STAT3 (CST 9145), anti-STAT3 (CST 4904), anti-phospho-ATF2 (CST 9221), anti-ATF2 (CST 9226), anti-phospho-IRF3 (CST 4947), anti-IRF3 (CST 4302), anti-PIMT (ab 70559), anti-FAM86A (sc 99414), anti-PRMT1 (CST 2449), anti-Tubulin (CST 2128), anti-GAPDH (sc 166545), anti-PTP1B (ABS40), anti-CSK (CST 4980), anti-GFP (sc 9996), anti-Flag (CST 8146), anti-HA (sc 7392), anti-acetylated Lysine (CST 2128), and anti-E-cadherin (CST 3195) antibodies were used. siRNA to human METTL18, PIMT, PRMT, and PKMT was purchased from Genolution (Seoul, Korea). AMI-1, pp2, stattic were purchased from Calbiochem (La Jolla, CA, USA). Phalloidin, S-adenosyl-L-methionine (SAM), lipopolysaccharide (LPS), cytochalasin B (Cyto B), VER-15508, and 17-AAG were purchased from Sigma Chemical Co. (St. Louis, MO, USA). siRNA sequences used in this study is listed in Supplementary Table 2.

#### Construction of expression vectors and virus production

A GFP-tagged wild-type actin construct (GFP-Actin-WT) and Myc-tagged wild type METTL18 construct (Myc-METTL18) were prepared via amplification with a typical culture method using competent *E. coli* (DH5α). The mutant constructs were prepared by the QuikChange Lightning Site-Directed Mutagenesis Kit. The pcDNA-HA, pcDNA-HA-tagged c-Src construct (Src-WT), Src Lys295 mutants (kinase-deficient HA-Src, Src-KD), and Src domain deletion mutants used in this study were the same as reported previously (<u>1</u>). All construct sequences were confirmed by the BigDye® Terminator v3.1 Cycle Sequencing Kit (Biosystems) from Macrogen, Inc (Seoul, Korea). The shRNA targeting region of the METTL18 (TRCN0000232974-shMETTL182974 and TRCN000023-2975-shMETTL182975) were obtained from Sigma (Supplementary Table 3). Lentiviruses containing shScramble, shMETTL182974, or shMETTL182975 were expressed by transient transfection using polyethylenimine (PEI, Sigma) into HEK293T cells along with pMD2.G (a gift from Yoon. K, Addgene 12259) and psPAX2 (a gift from Yoon. K, Addgene 12260), and the viral supernatants were collected after 48 h.

## Supplementary Figure Legends

**Supplementary Figure S1.**
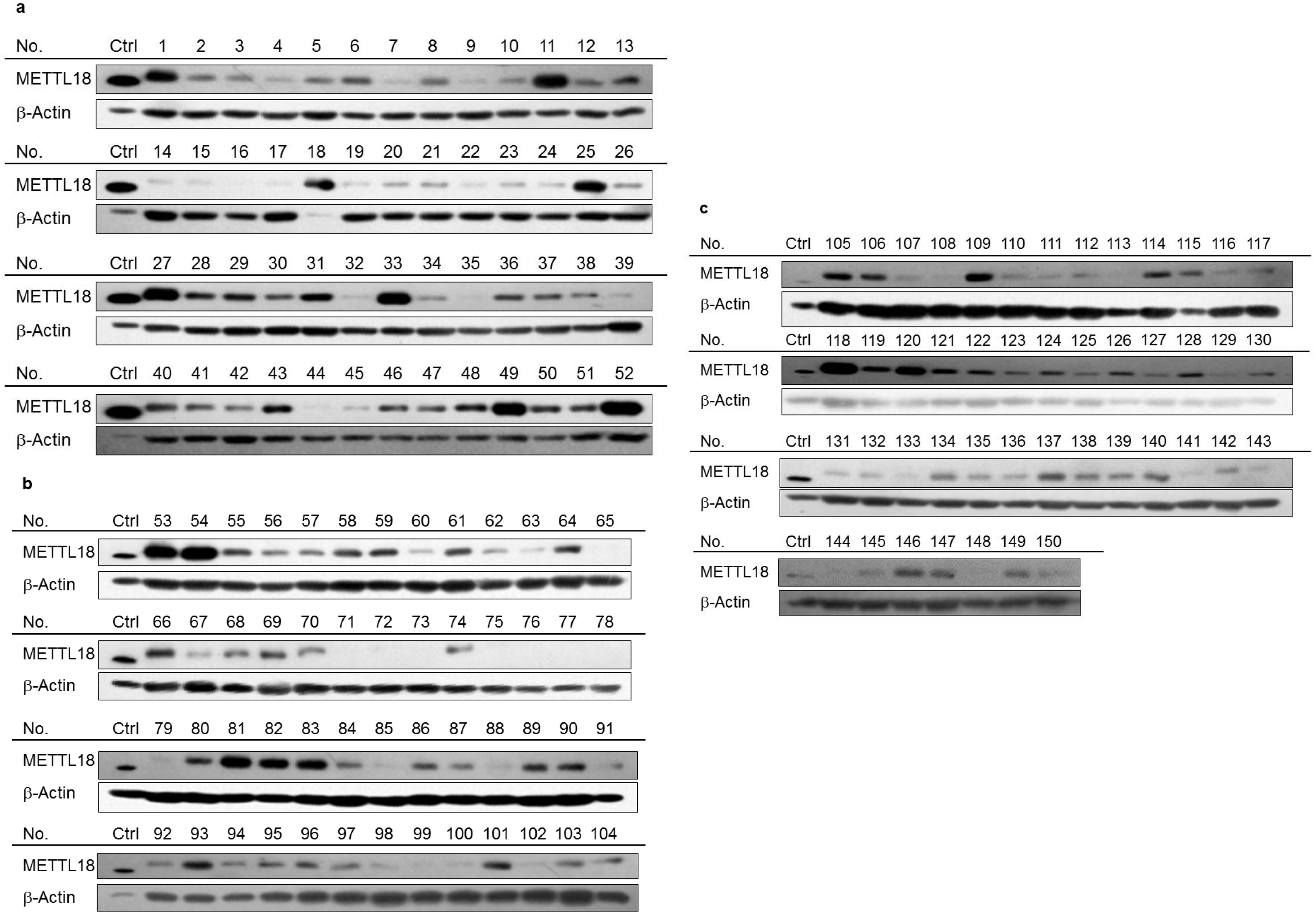
Protein levels of METTL18 in breast cancer tissues from patients. **(A-C)** Protein level of PHMT was determined by Western blotting analysis from breast cancer tissues of patients. Breast cancer tissues of patients were acquired from Samsung Medical Center (Seoul, Korea).

**Supplementary Figure S2.**
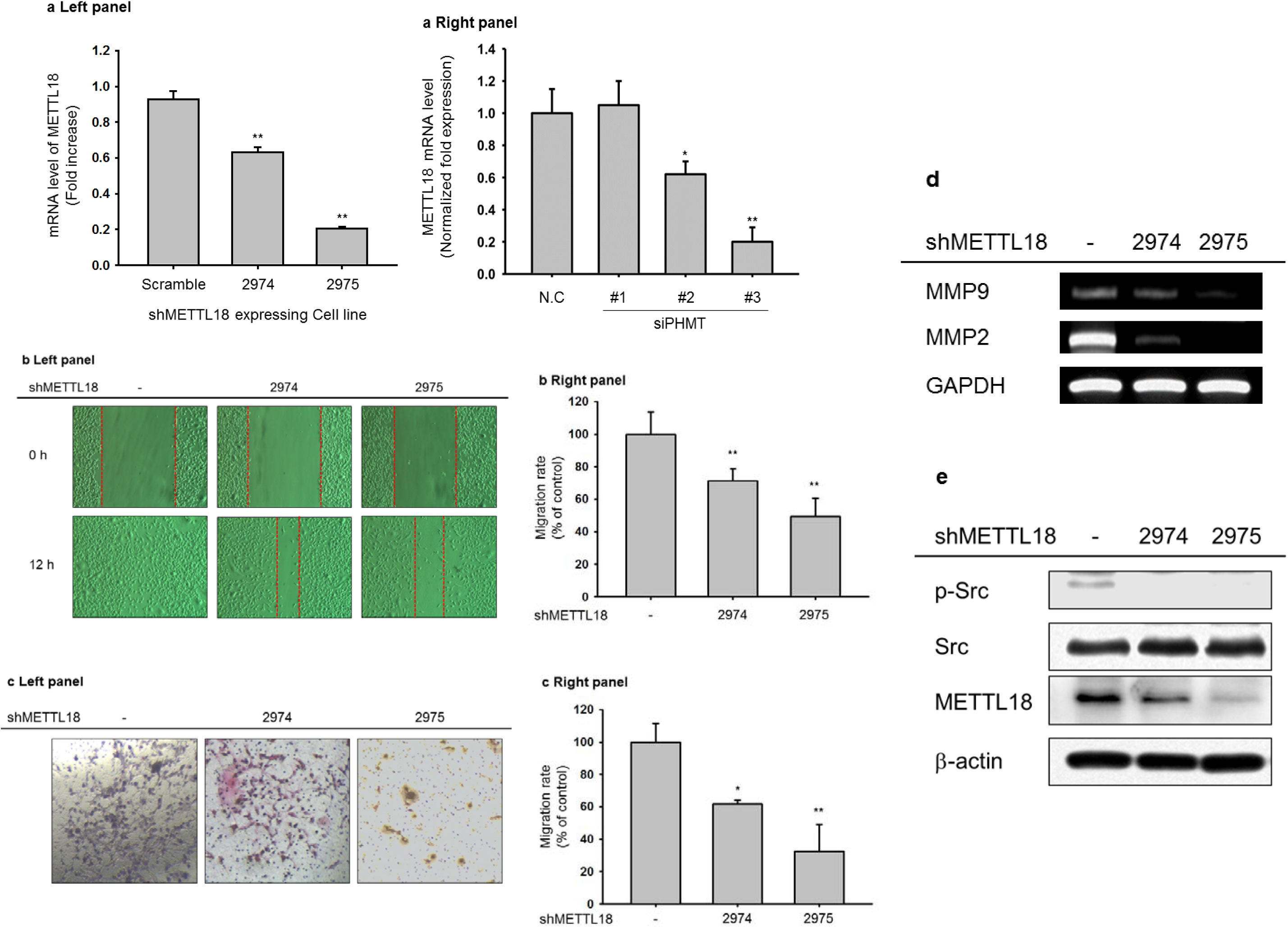
The mRNA expression levels of METTL18 from MDA-MB-231 cells expressing shRNA to METTL18. (A Left panel) shMETTL18-expressing MDA-MB-231 cell lines were established by lentiviral infection. Different types of siRNA to METTL18 were also transfected to MDA-MB-231 cells (A Right panel). The knockdown level of METTL18 from these cells was determined by real-time PCR. (B and C) *In vitro* migration and invasion assays were performed with MDA-BM-231 cells expressing shMETTL18 (#2974 or #2975). (D) Expression levels of MMP9 and MMP2 in shMETTL18-expressing MDA-MB-231 cell were determined by RT-PCR (E) Phospho-Src and METTL18 levels were detected by Western blotting analysis. N.C.: negative control. *: p<0.05 and **: p<0.01 compared to shScramble group or N.C.

**Supplementary Figure S3.**
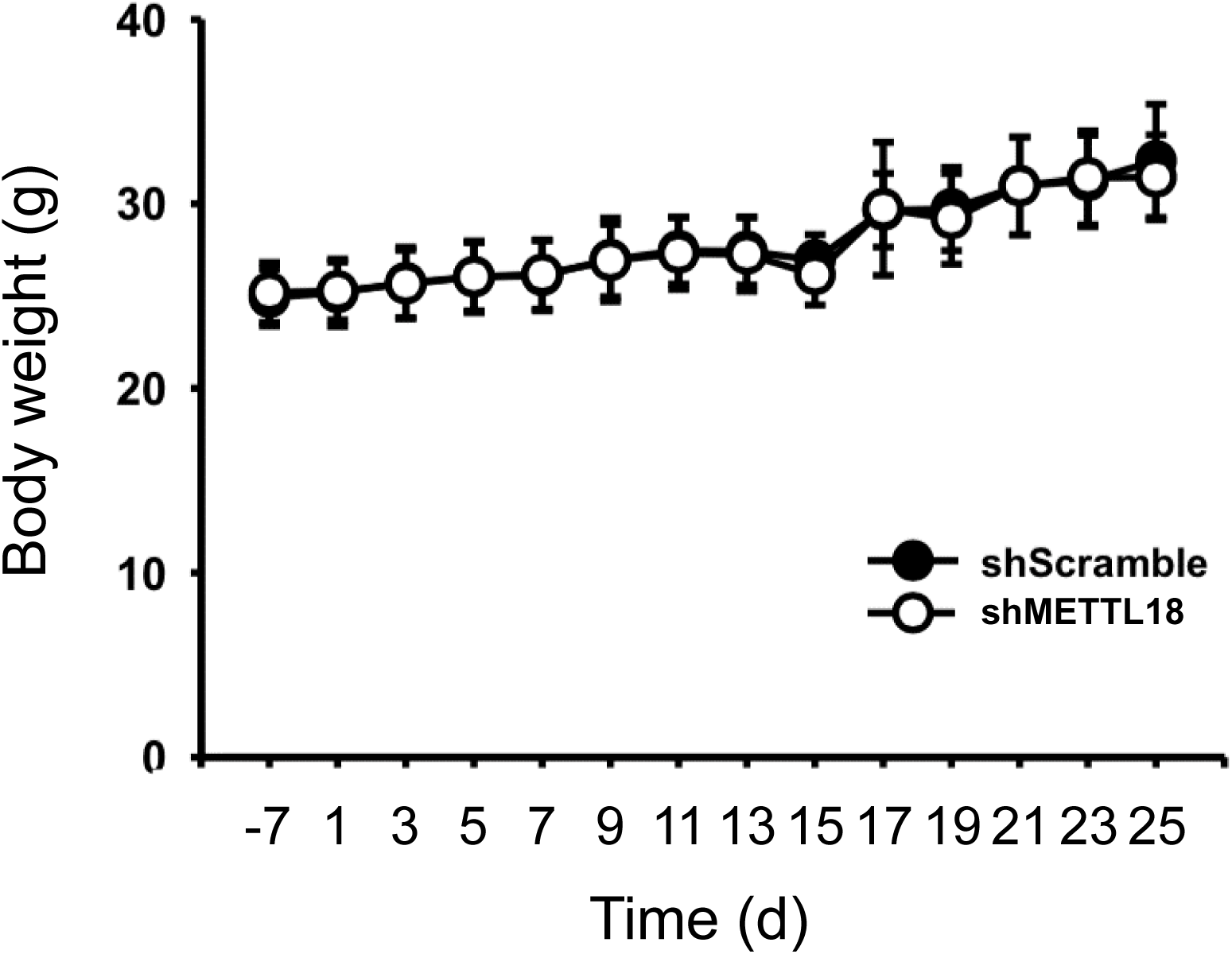
Body weight profile of mice injected with shScramble or shMETTL18 (shPHMT)-expressing MDA-MB-231 cells. Body weight of mice in each group was measured with a digital balancer every day.

**Supplementary Figure S4.**
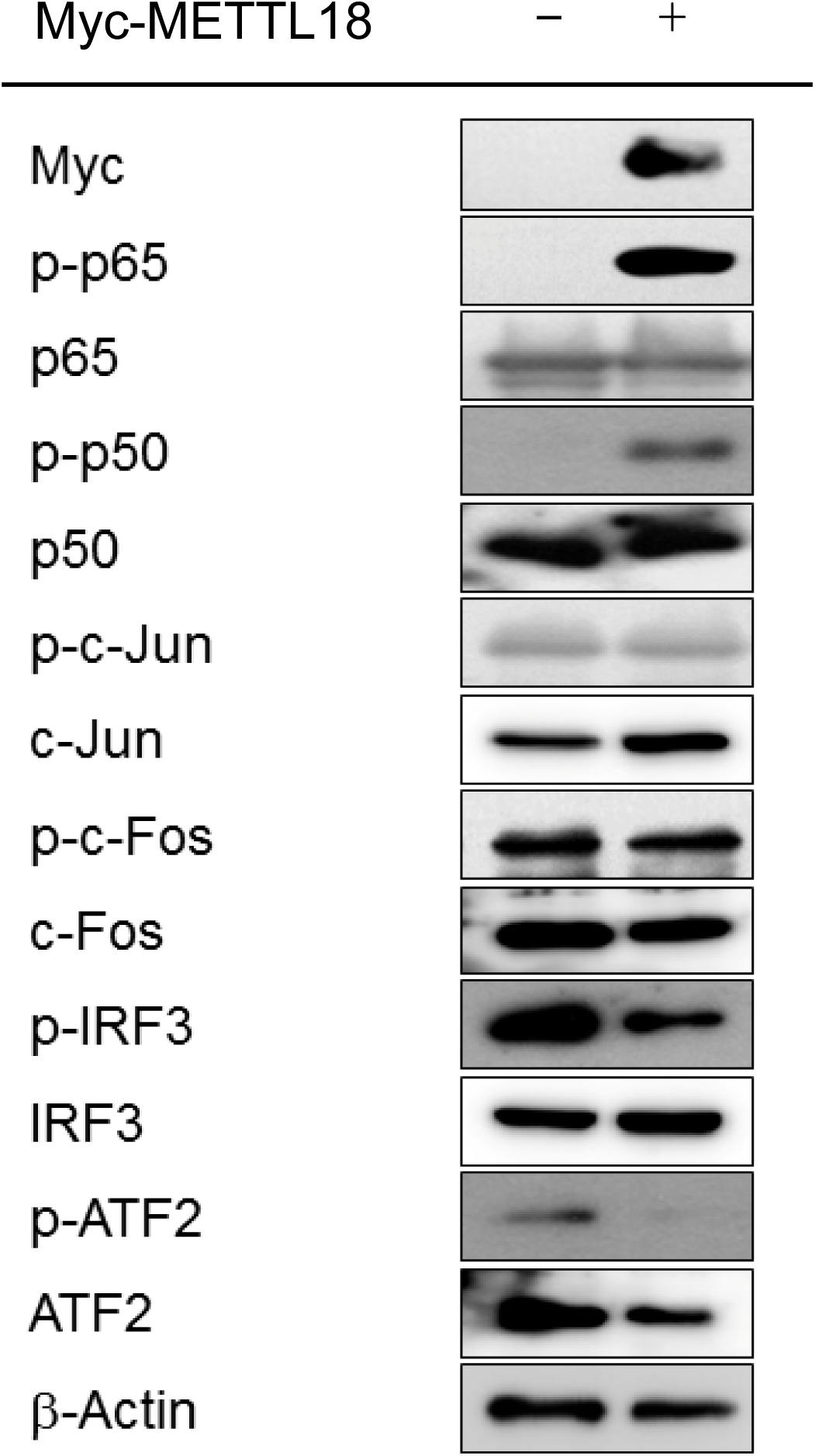
Protein levels of transcription factors involved in tumorigenic responses from Myc-METTL18-overexpressing MDA-MB-231 cells. The total and phospho-protein levels of p65, p50, c-Jun, c-Fos, IRF3, ATF2, Myc and β-actin were determined by Western blotting analysis.

**Supplementary Figure S5.**
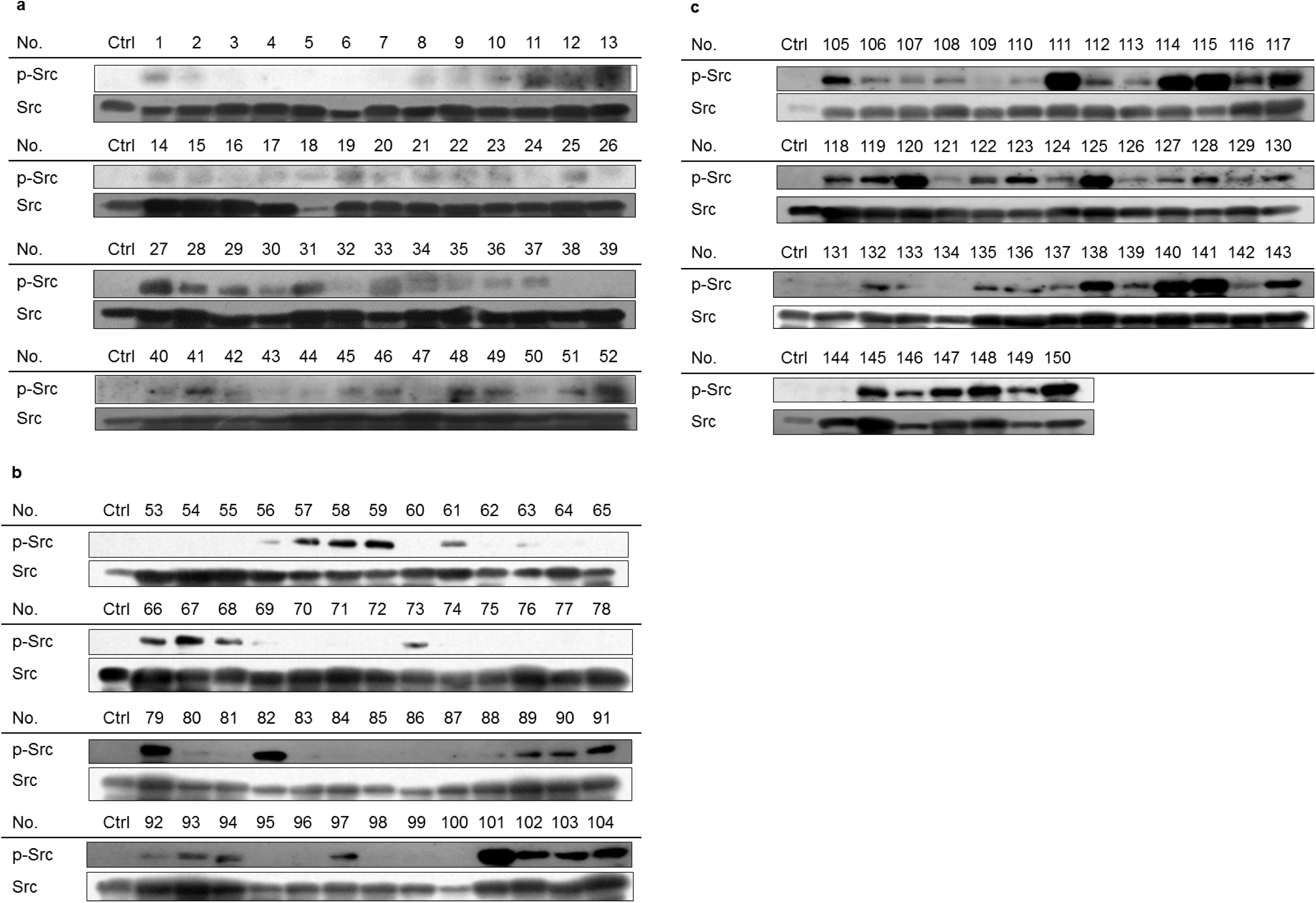
Phospho-Src level in breast cancer tissues of patients. **(A-C)** The total and phospho-protein levels of Src was determined by Western blotting analysis from breast cancer tissues of patients. Breast cancer tissues of patients was acquired from Samsung Medical Center (Seoul, Korea).

**Supplementary Figure S6.**
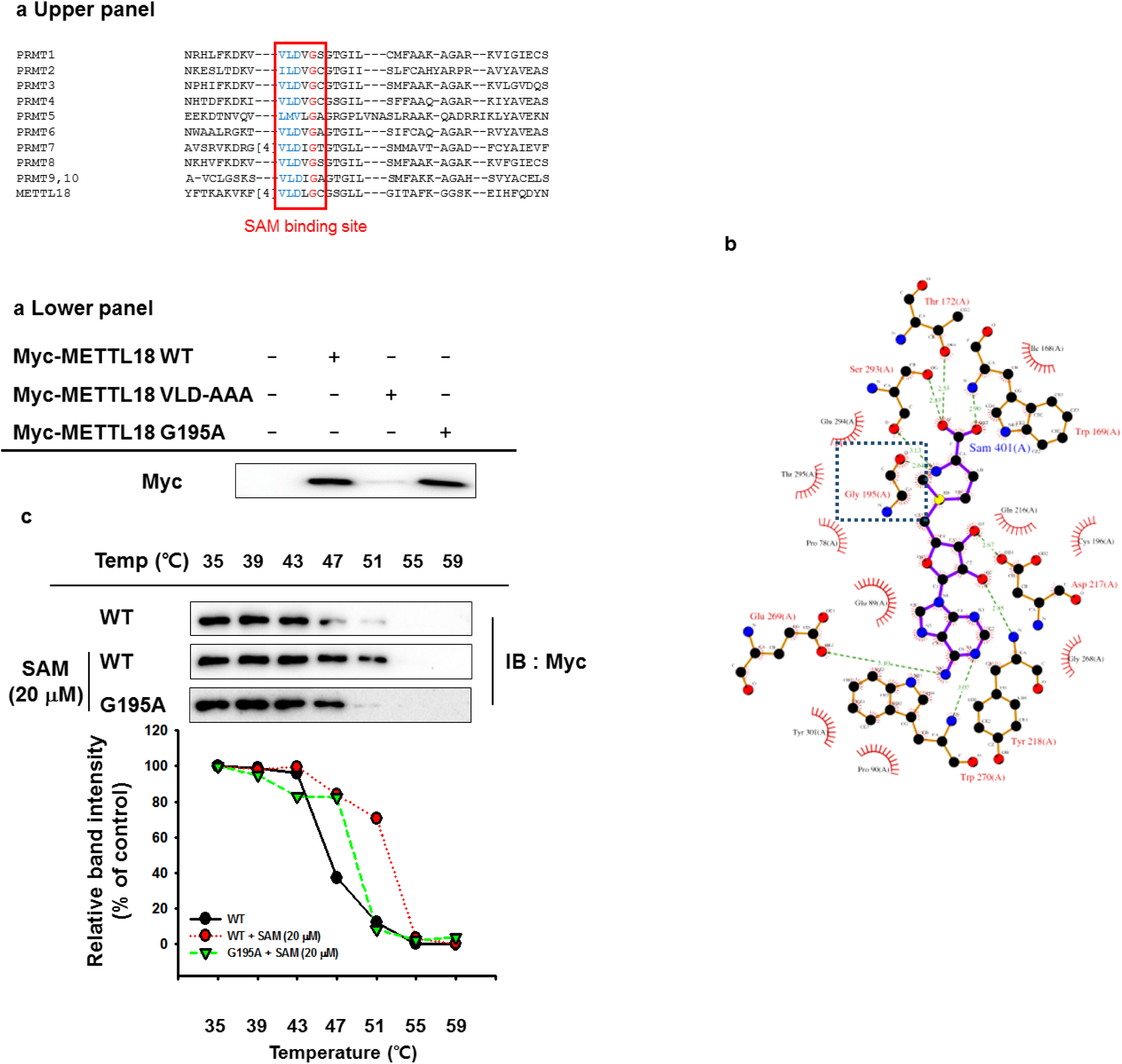
Expected SAM binding site of METTL18. **(A Upper panel)** METTL18 amino acid sequences were aligned with PRMT1-10 using Clustal Omega program. **(A Lower panel)** Protein expression levels of METTL18 mutants were determine by Western blotting analysis. **(B)** Putative binding site of SAM in METTL18 was speculated by PDB (PDB ID : 4rfq, Tempel W, Ravichandran M, Li Y, Walker JR, Bountra C, Arrowsmith CH, Edwards AM, Brown PJ, Hong BS, Structural Genomics Consortium). **(C)** A cellular thermal shift assay was carried out to check the thermal stability of METTL18 or METTL18 G195A in the presence and absence of SAM. Western blotting level of METTL18 at 35 °C was considered to be 100%.

**Supplementary Figure S7.**
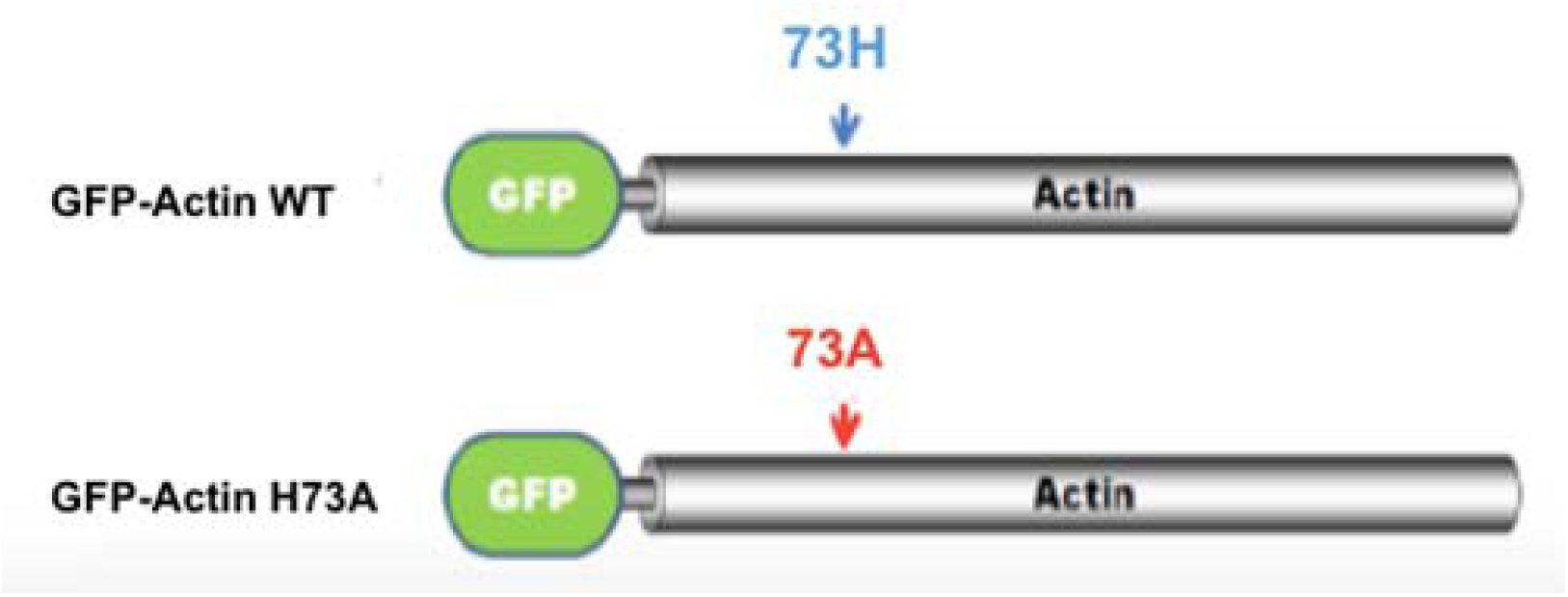
Schematic diagram of GFP-Actin-wild type (WT) and its mutant at 73-histidine to alanine residue. Putative histidine methylation site at 73 amino acid residue is mutated to alanine.

**Supplementary Figure S8.**
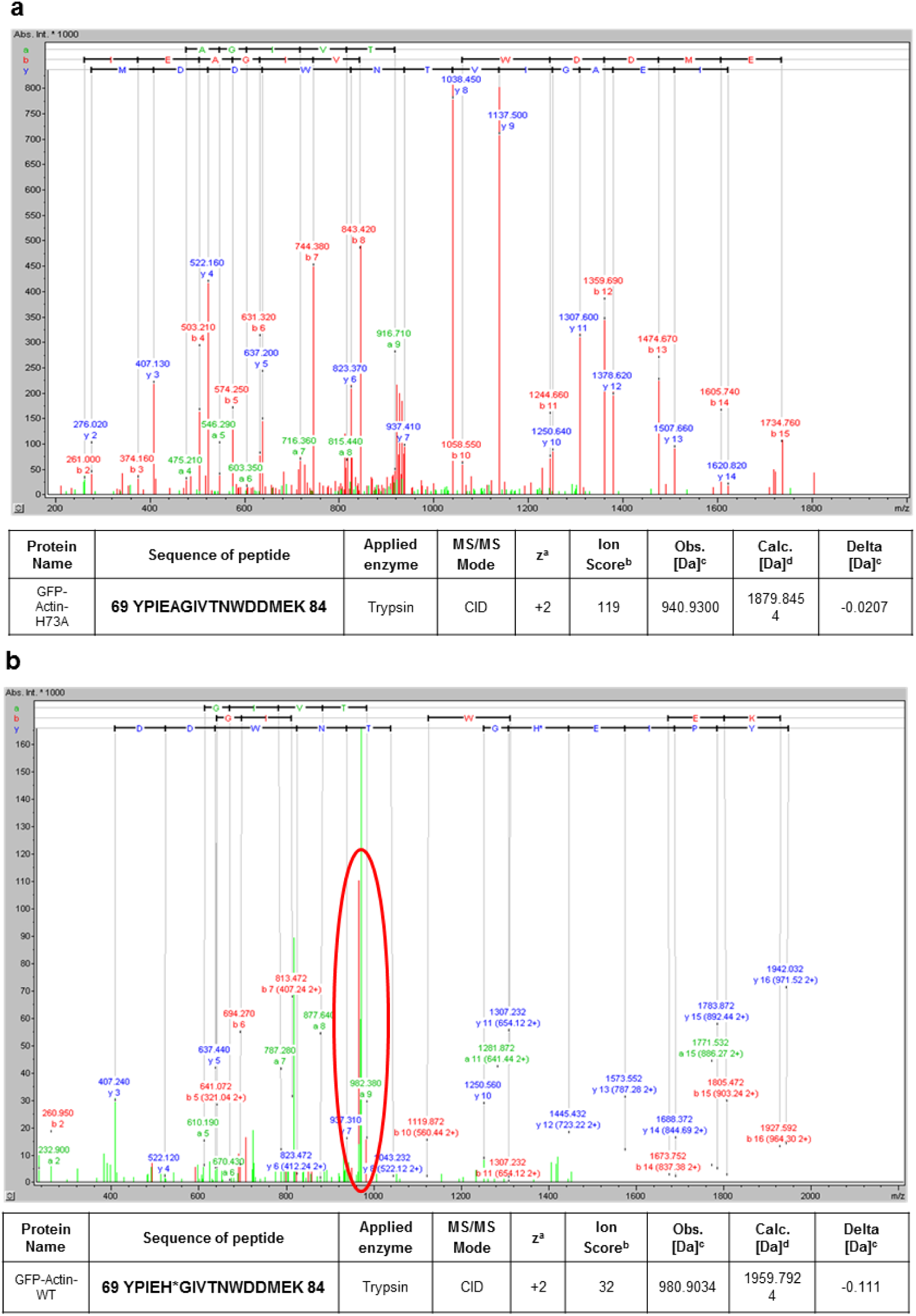
Analysis of histidine methylation site in actin by Mass spectrometry. **(A and B)** Histidine methylation of actin at 73-histidine residue by METTL18 was examined by comparison with actin (B) and its mutant at 73-histidine (A) by mass spectrometric analysis.

**Supplementary Figure S9.**
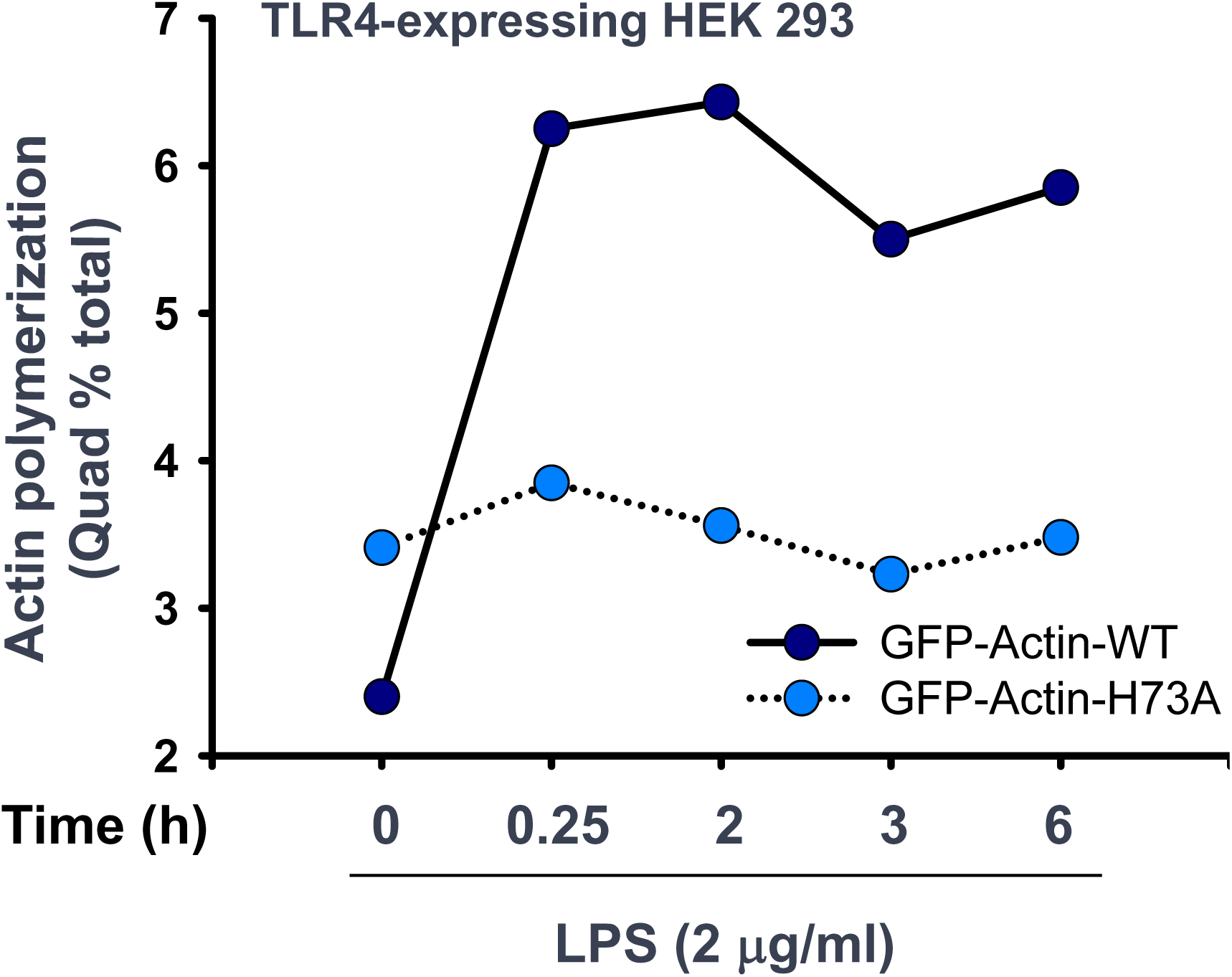
Role of 73-histidine residue in LPS-induced actin polymerization. TLR4-expressing HEK293 cells transfected with GFP-Actin-WT or GFP-Actin-H73A were activated in the presence or absence of LPS (2 μg/ml). After staining with actin phalloidin, fluorescence of the cells was analyzed by flowcytometry.

**Supplementary Figure S10.**
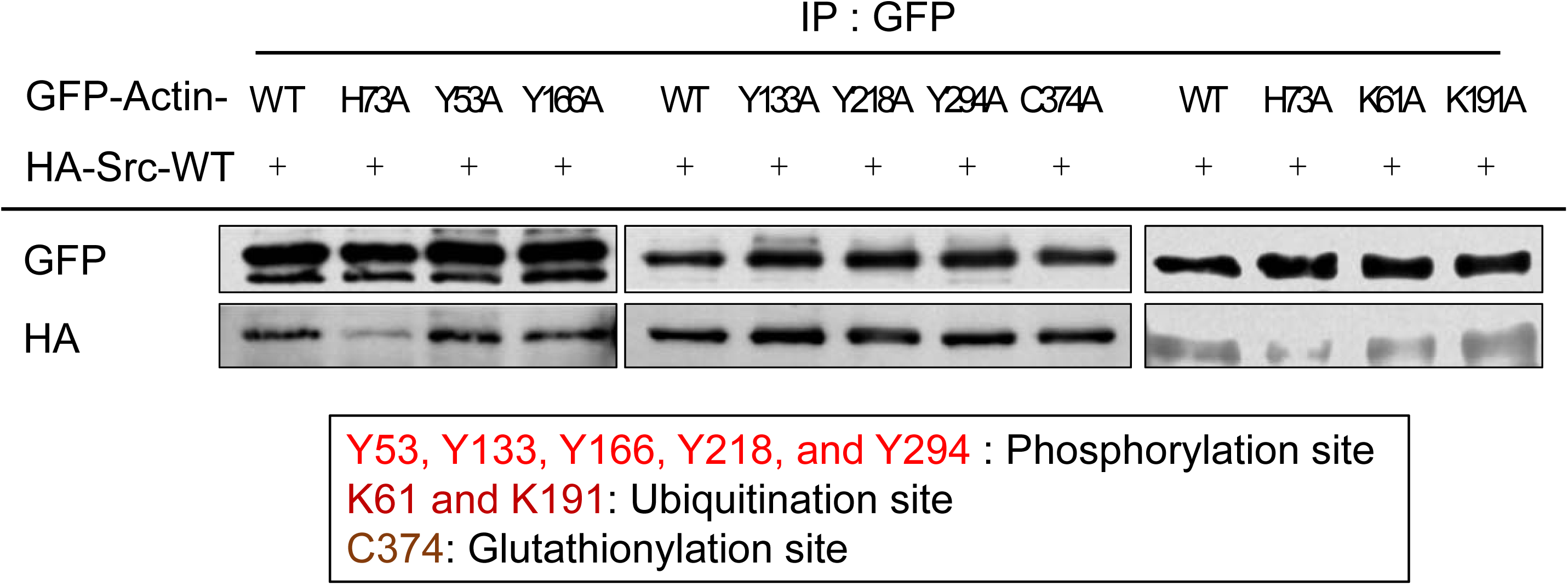
Effect of post-translational modification of actin at histidine, tyrosine, cysteine or lysine residue on binding between Src and actin. Src/actin binding level was confirmed by immunoprecipitation and Western blotting analysis with GFP-Actin mutants at tyrosine phosphorylation sites (Y133, Y218, and Y294), histidine methylation site (H73), ubiquitination sites (K61 and K191), and glutathionylation site (C374).

**Supplementary Figure S11.**
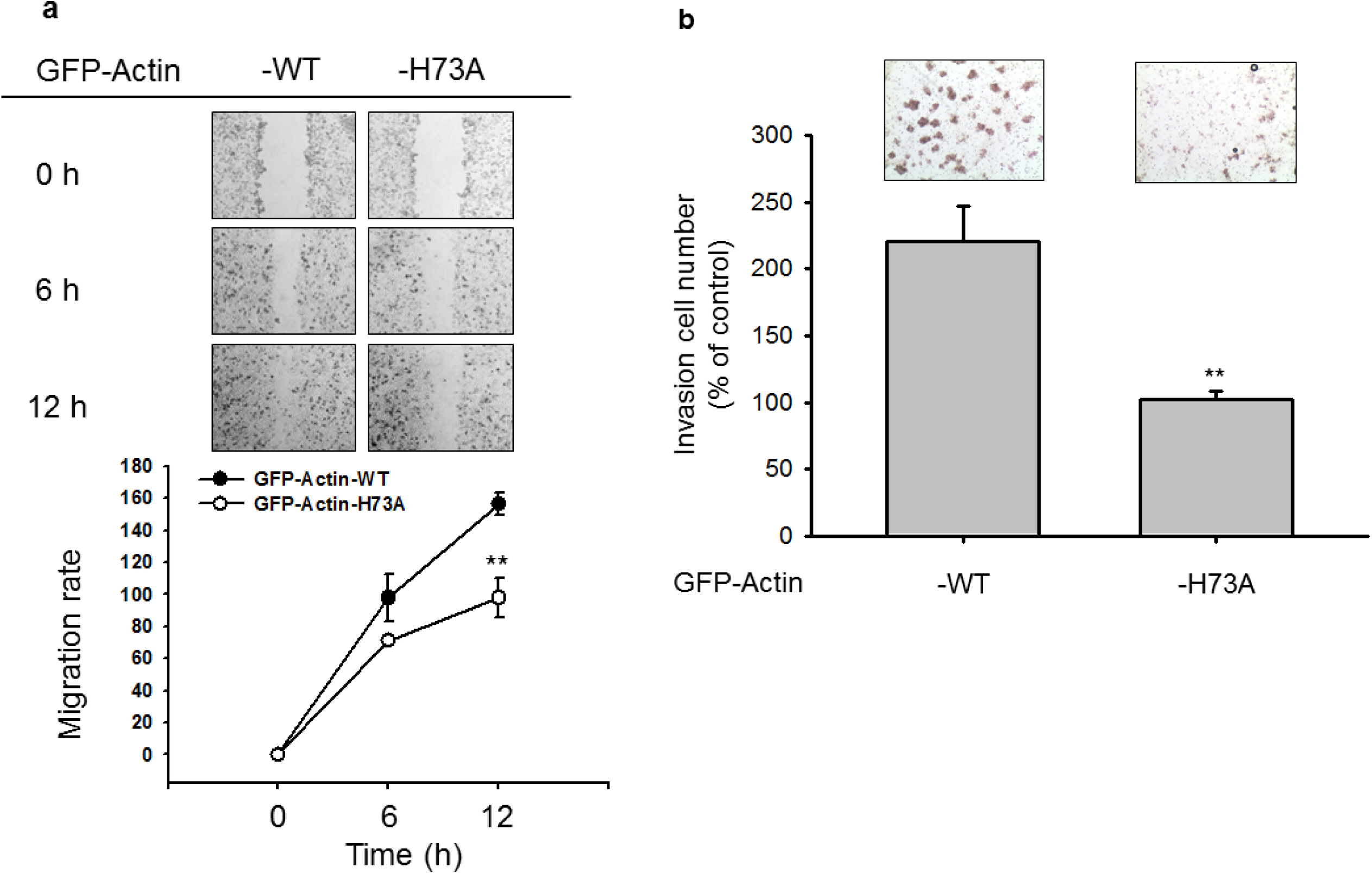
Role of histidine methylation at 73-histidine on the migration and invasion of MDA-MB-231 cells. **(A and B)** Migration and invasion patterns of MDA-MB-231 cells transfected with GFP-Actin-WT or –H73A were examined by wound healing (A) and invasion (B) assay. **: p<0.01 compared to wild type.

**Supplementary Figure S12.**
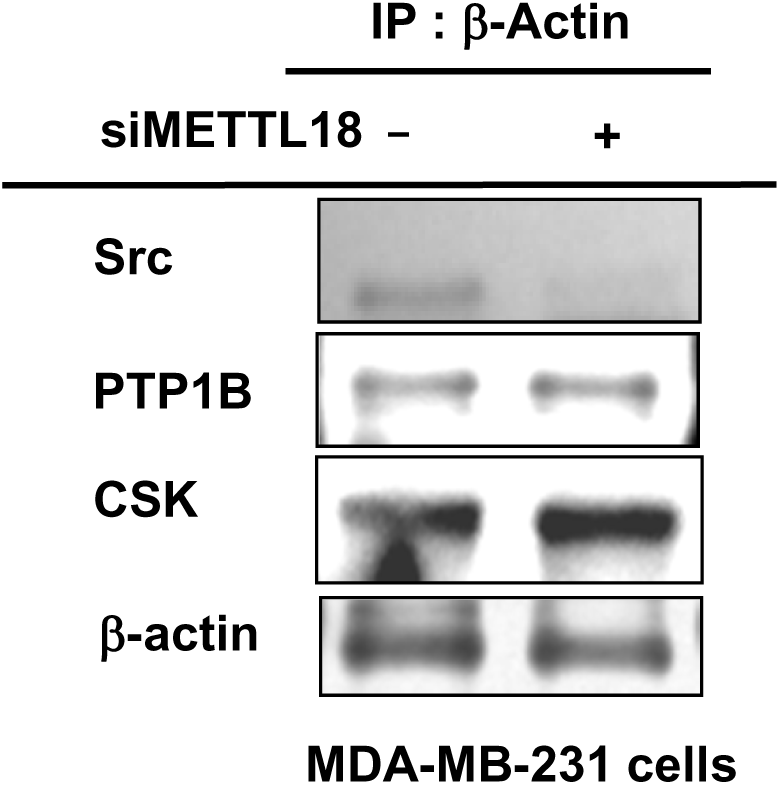
Role of METTL18 on the binding patterns between actin and METTL18. Critical role of METTL18 in controlling molecular interaction between actin and Src was analyzed by immunoprecipitation and Western blotting analysis from siMETTL18-transfected MDA-MB-231 cells. The protein levels of Src, PTP1B, CSK, and β-actin were analyzed from the IP complex.

**Supplementary Figure S13.**
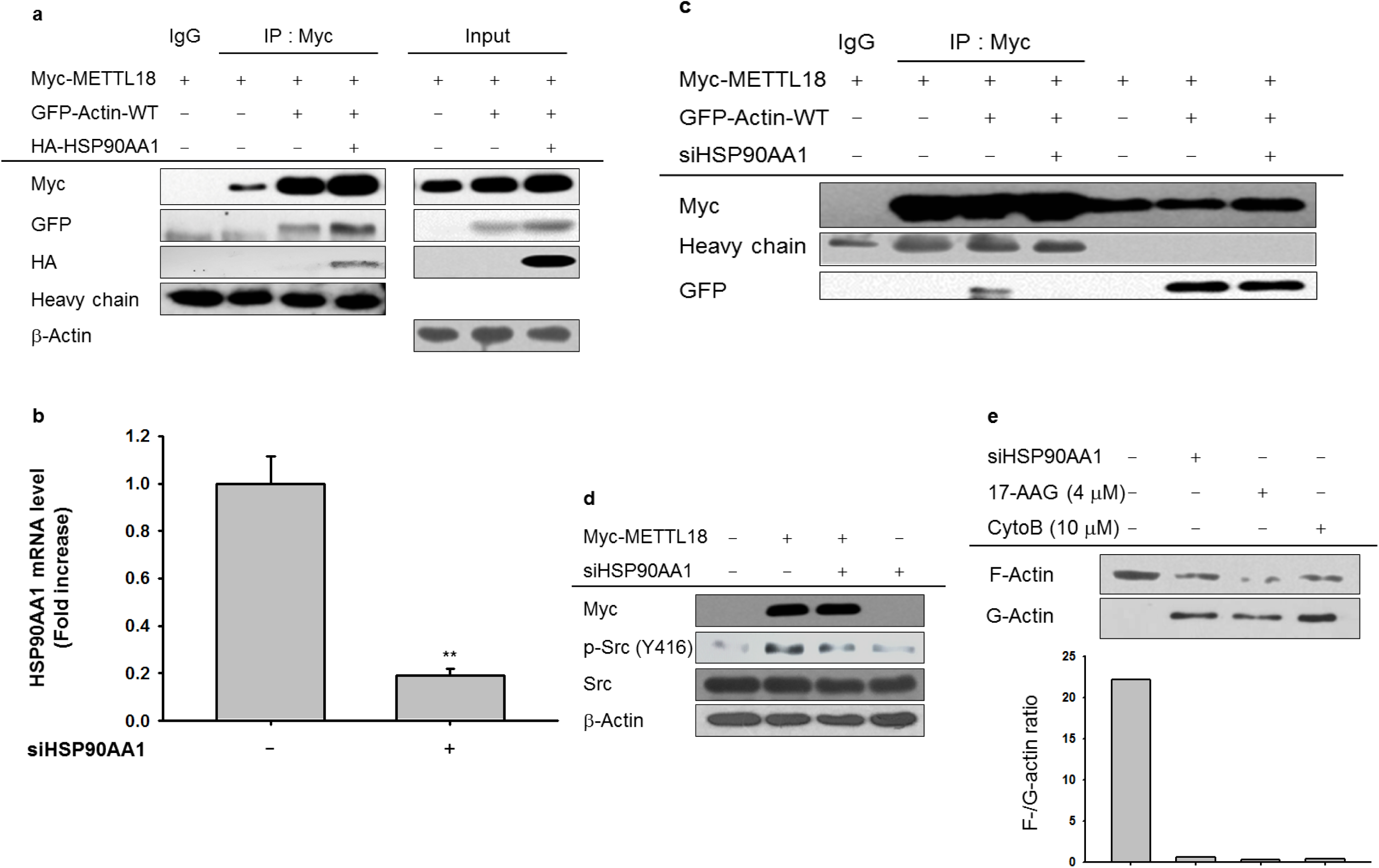
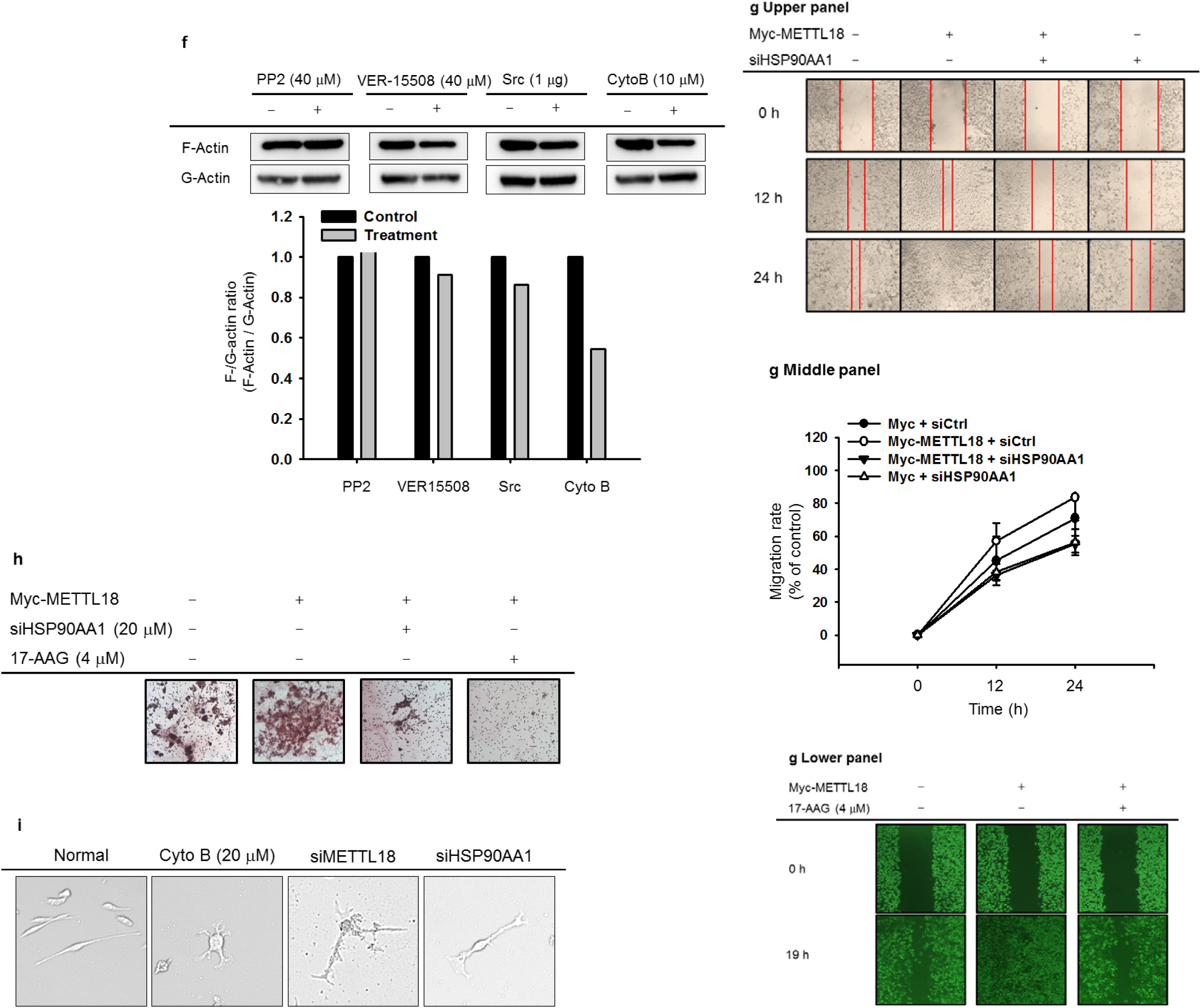
Role of HSP90AA1 on the regulation of PHMT/actin binding and Src activation. **(A)** Binding level of HSP90AA1 to PHMT and actin was detected by immunoprecipitation and Western blotting analysis. **(B)** Knockdown level of HSP90AA1 mRNA under siHSP90AA1 treatment condition in MDA-MB-231 cells was determined by real-time PCR. **(C)** Role of HSP90AA1 on PHMT/actin interaction was assessed by detection of GFP-actin level by immunoprecipitation with Myc-PHMT and Western blotting analysis from siHSP90AA1-treated MDA-MB-231 cells. **(D)** Effect of siHSP90AA1 on PHMT-induced Src phosphorylation was determined by analyzing p-Src level from Myc-PHMT-overexpressed MDA-MB-231 cells. **(E and F)** The role of HSP90AA1 on the stimulation of actin polymerization was determined by an F-actin/G-actin assay in MDA-MB-231 cells transfected with siHSP90AA1, 17-AAG (4 μM), CytoB (10 μM), PP2 (40 μM), VER155508 (40 μM), or Src (1 μg). **(G, H, and I)** Effect of siHSP90AA1 or 17-AAG on migration **(G)**, invasion **(H)** and morphological change **(I)** of MDA-MB-231 cells transfected with Myc-PHMT. **: p<0.01 compared to normal.

**Supplementary Figure S14.**
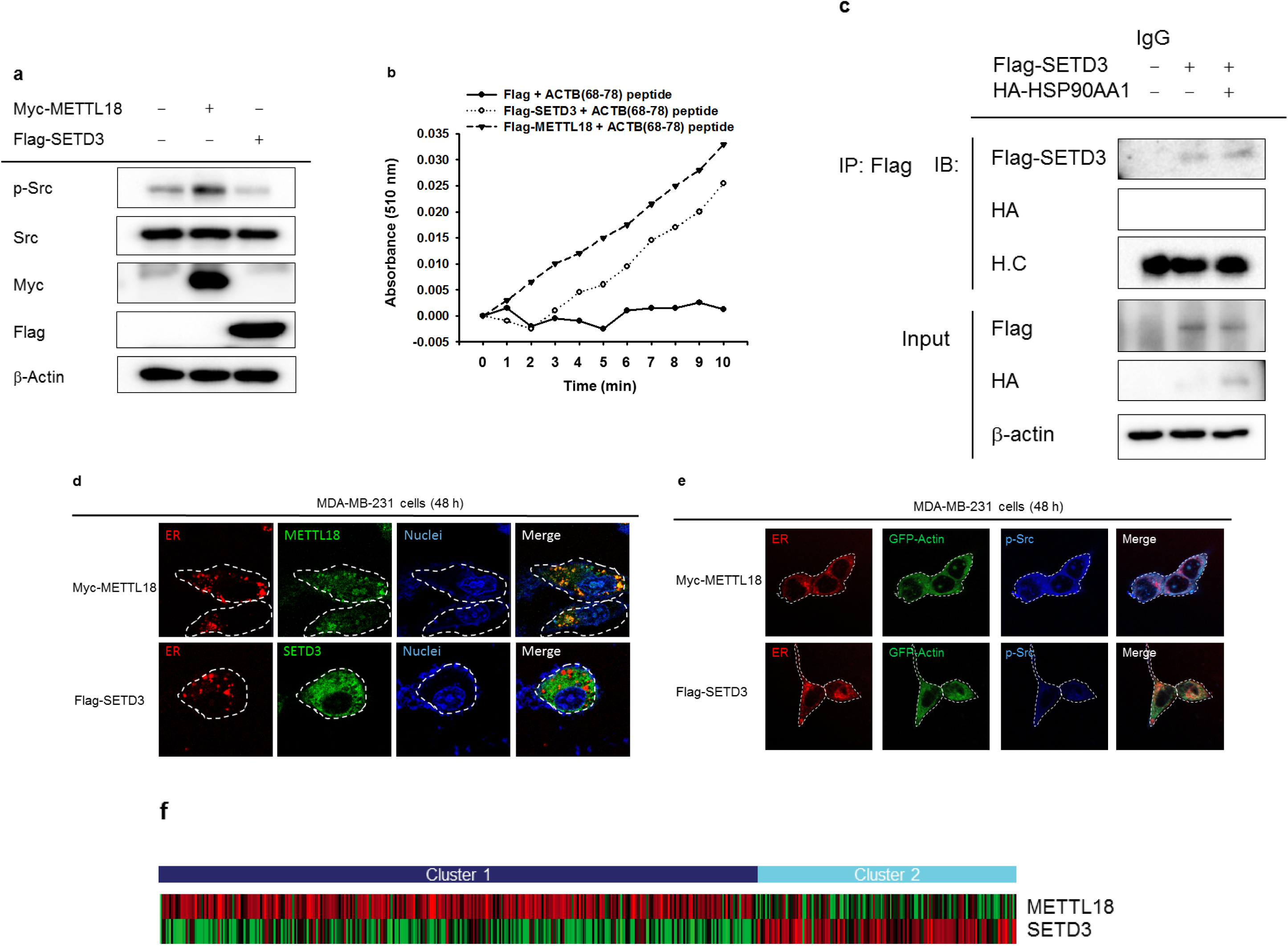
Distinctive feature of SETD3 or METTL18 in Src activation, actin-derived peptide methylation, and expression pattern in breast tumor patients. **(A)** Phosphorylation of Src was examined by Western blotting analysis in METTL18- or SETD3-overexpressed MDA-MB-231 cells. **(B)** Histidine-methylating activity of SETD3 or METTL18 in histidine residue of ACTB(68-78) peptide derived from actin was demonstrated with colorimetric SAM510 methyltransferase assay kit. **(C)** Binding level between SETD3 and HSP90 was determined by immunoblotting and Western blotting analysis from MDA-MB-231 cells transfected with Flag-SETD3 and HA-HSP90AA1. **(D and E)** Cellular distribution of METTL18 or SETD3 in MDA-MB-231 cells transfected with Myc-METTL18 or Flag-SETD3 was detected by Confocal microscopic analysis. **(F)** Gene expression patterns of SETD3 and METTL18 in breast cancer tumor patients were analyzed with heatmap. Samples were obtained from TCGA dataset.

## Supplementary Tables

**Supplementary Table 1.**
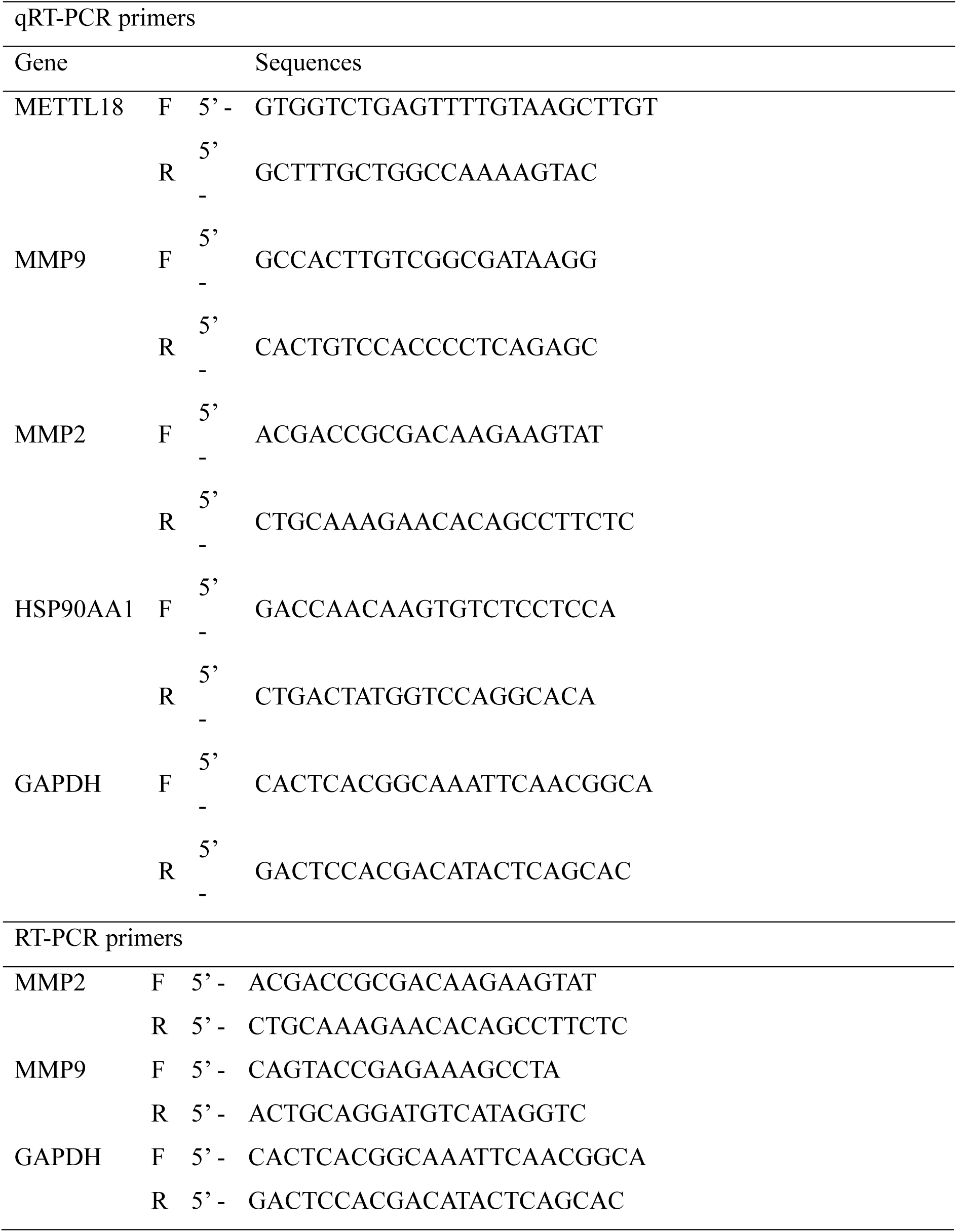
List of primers used for mRNA analysis by real-time PCR and RT-PCR

**Supplementary Table 2.**
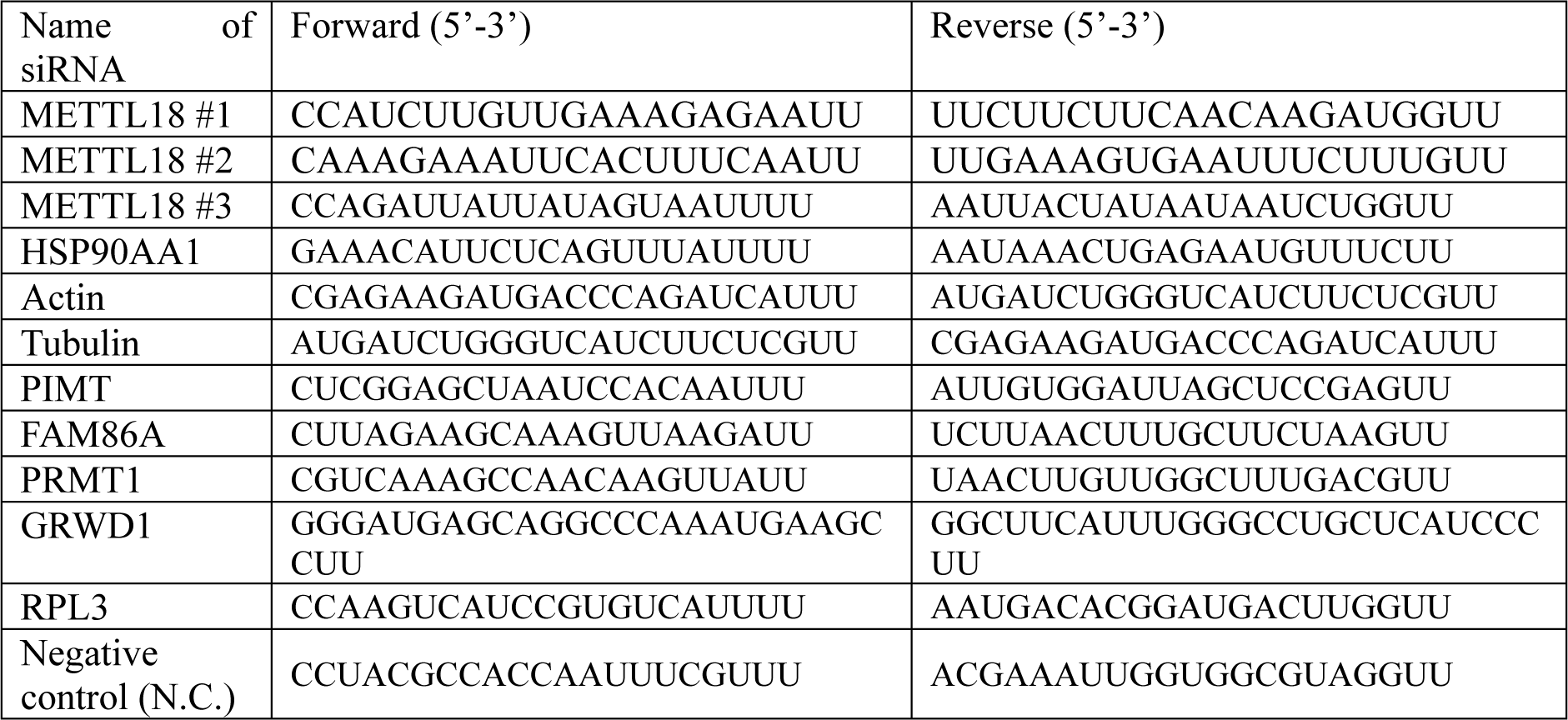
List of primers to prepare siRNA to METTL18, HSP90AA1, actin, and tubulin.

**Supplementary Table 3.**
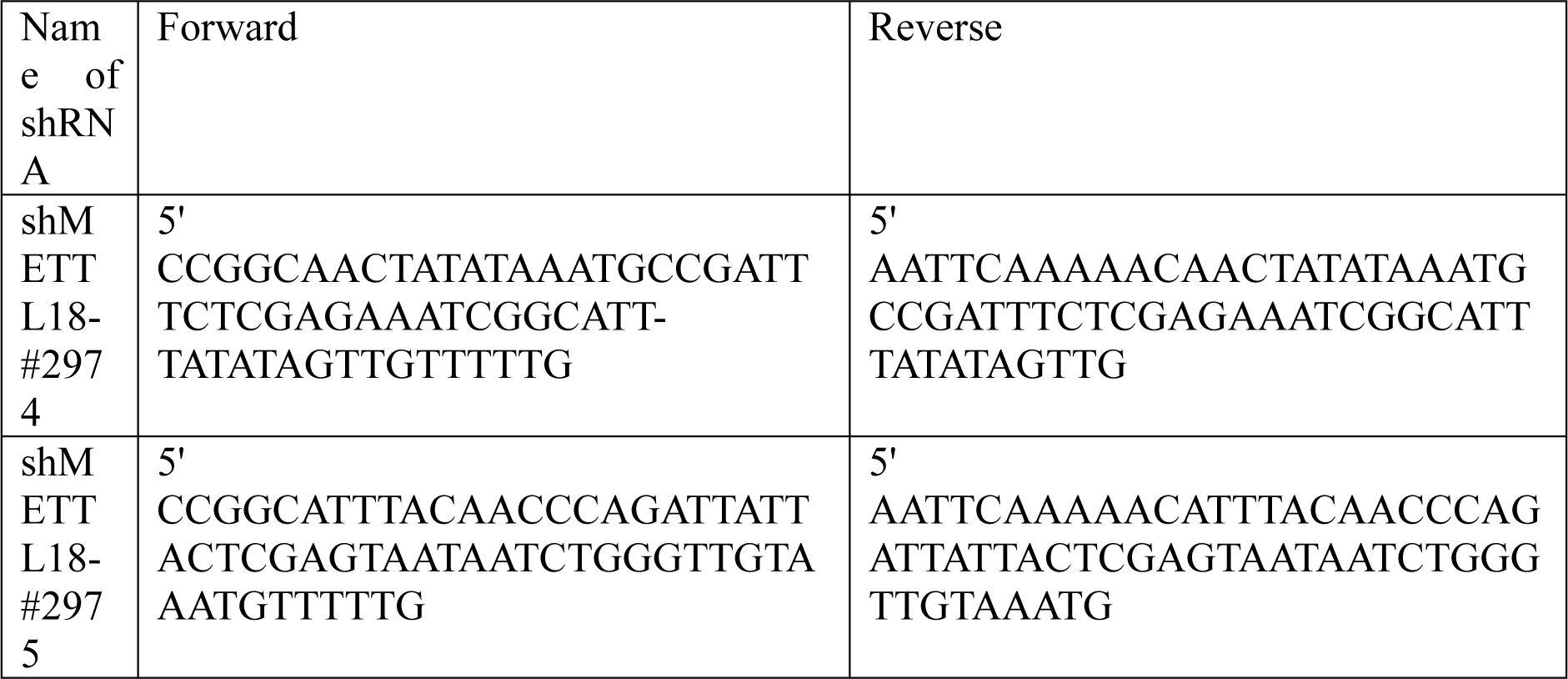
List of primers to prepare METTL18 shRNA expressing constructs.

